# LINE-1 retrotransposon activation intrinsic to interneuron development

**DOI:** 10.1101/2022.03.20.485017

**Authors:** Gabriela O. Bodea, Juan M. Botto, Maria E. Ferreiro, Francisco J. Sanchez-Luque, Jose de los Rios Barreda, Jay Rasmussen, Muhammed A. Rahman, Laura R. Fenlon, Natasha Jansz, Carolina Gubert, Patricia Gerdes, Liviu-Gabriel Bodea, Prabha Ajjikuttira, Darwin J. Da Costa Guevara, Linda Cumner, Charles C. Bell, Peter Kozulin, Victor Billon, Santiago Morell, Marie-Jeanne H.C. Kempen, Chloe J. Love, Karabi Saha, Lucy M. Palmer, Adam D. Ewing, Dhanisha J. Jhaveri, Sandra R. Richardson, Anthony J. Hannan, Geoffrey J. Faulkner

## Abstract

Retrotransposons are a reservoir of cis-regulatory innovation^1–3^. Developmental programs that activate these elements could, in principle, manifest in lineage-specific retrotransposition. Somatic LINE-1 (L1) retrotransposon insertions have been detected in human and non-human primate neurons^4–7^. It is however unknown whether L1 is mobile in only some neuronal lineages, or therein regulates neurodevelopmental genes. Here, we report programmed L1 activation by SOX6, a transcription factor critical for parvalbumin (PV) interneuron development^8–10^. PV^+^ neurons permit L1 mobilization *in vitro* and *in vivo*, harbor unmethylated L1 promoters, and express full-length L1 mRNAs and proteins. Via nanopore long-read sequencing, we identify unmethylated L1 promoters proximal to PV^+^ neuron genes. One such L1, which promotes transcription of a novel CAPS2 gene isoform, significantly enhances neuron morphological complexity when phenotyped *in vitro*. These data highlight the contribution made by L1 cis-regulatory elements to PV^+^ neuron development and transcriptome diversity, uncovered due to L1 mobility in this milieu.

## Maintext

The retrotransposon LINE-1 (L1) comprises ∼18% of the human and mouse genomes, and is a notable source of protein-coding gene and cis-regulatory variation^11–13^. To mobilize, L1 initiates transcription of a full-length (>6kbp) mRNA from its internal 5ʹUTR promoter. The mRNA encodes two proteins, denoted ORF1p and ORF2p, that mediate L1 retrotransposition^14–16^. Heritable L1 insertions principally arise in the early embryo and primordial germ cells^17–21^. Somatic L1 retrotransposition can occur in the brain, as first revealed by L1-EGFP retrotransposition reporter experiments^22–24^ and later confirmed by single-neuron (NeuN^+^) genomic analyses^4–7^. However, the key question remains of whether specific neuronal lineages are more permissive for somatic retrotransposition than others.

To resolve the spatial and cell type specificity of somatic L1 mobility *in vivo*, we generated a transgenic L1-EGFP mouse line (**Fig. 1a** and **Extended Data Fig. 1a**) harboring a retrotransposition reporter based on L1.3, a highly mobile human L1 element^25,26^. L1.3 was expressed from its native promoter and incorporated T7 and 3×FLAG epitope tags on ORF1p and ORF2p, respectively (**Fig. 1a**). Immunofluorescence revealed EGFP^+^ neurons (**Fig. 1b**). In agreement with prior experiments^22^, nearly all EGFP^+^ cells were found in the brain, apart from occasional EGFP^+^ ovarian interstitial cells (**Extended Data Fig. 1b-e**). Tagged ORF1p and ORF2p expression was observed in EGFP^+^ neurons, indicating the L1 protein machinery coincided with retrotransposition (**Fig. 1c**,**d**). Guided by their location and morphology, we hypothesized the EGFP^+^ cells were predominantly parvalbumin (PV) inhibitory interneurons. Immunostaining indicated 85.4% of EGFP^+^ hippocampal cells were PV^+^ neurons, on average (**Fig. 1e**,**f**). PV^+^/EGFP^+^ neurons were found throughout the hippocampal dentate gyrus (DG) and cornu ammonis regions 1-3 (CA1-3, referred to here as CA) (**Fig. 1g**) but were infrequent in the cortex (**Fig. 1h**). EGFP^+^ cells also expressed GAD1 (**Extended Data Fig. 1f**), another inhibitory interneuron marker^27^. L1-EGFP retrotransposition was thus most common in the PV^+^ neuron lineage.

**Fig. 1:**
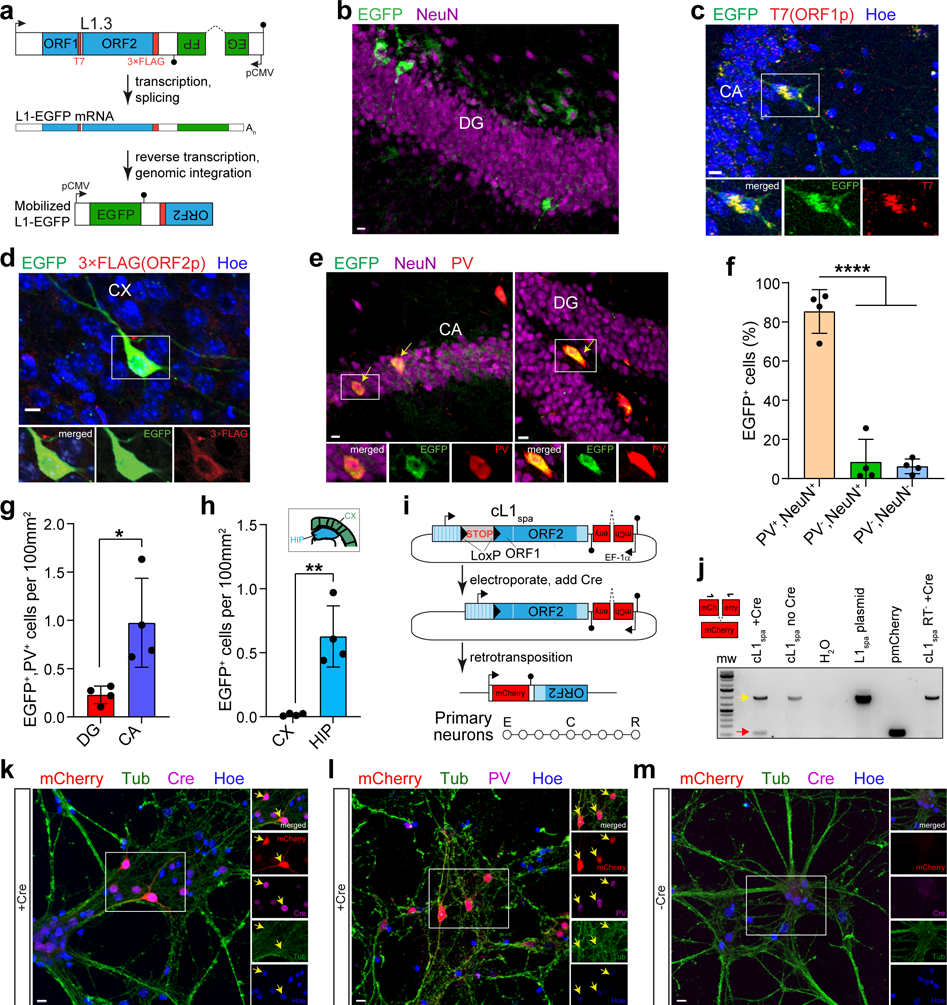
L1 retrotransposition in PV^+^ neurons. **a,** L1-EGFP reporter schematic. A mobile human L1 (L1.3) is expressed from its native promoter, harbors epitope tagged ORF1 (T7) and ORF2 (3×FLAG) sequences, and carries an EGFP indicator cassette. The EGFP is antisense to the L1, incorporates a ɣ-globin intron in the same orientation as the L1, and terminated by a polyadenylation signal (filled black lollipop). L1-EGFP retrotransposition removes the ɣ-globin intron, enabling EGFP expression. **b,** Example EGFP^+^ cells detected in the hippocampus. DG: dentate gyrus. **c,** Representative confocal image of ORF1p (T7) immunostaining of L1-EGFP adult mouse brain. Image insets show a selected cell in merged and single channels for EGFP and ORF1p. CA: cornu ammonis. **d,** As for (c), except for ORF2p (3×FLAG) in cortex (CX). **e,** EGFP and PV immunostaining of L1-EGFP mouse coronal hippocampus sections. Yellow arrows indicate EGFP^+^ neurons. **f,** Percentage of hippocampal EGFP^+^ cells colocalized with NeuN and PV. \*\*\*\**P*=0.0001, one-way ANOVA with Tukey’s post-hoc test, *N*(mice)=4. **g,** Distribution of EGFP^+^/PV^+^ cells counted in the main hippocampal substructures. \**P*=0.02, two-tailed t test. **h,** EGFP^+^ cell counts in cortex and hippocampus (HIP). \*\**P*=0.002, two-tailed t test. Note: panels (f-h) represent data as mean ± SD. **i,** cL1_spa_ reporter schematic. A mobile mouse L1 (L1_spa_) is expressed from its native monomeric 5′UTR, harbors a LoxP-Stop-LoxP cassette, and carries an mCherry indicator cassette. Cre-Lox recombination removes the Stop, enabling L1_spa_ transcription and retrotransposition. Primary neurons were electroporated (E) with cL1_spa_, transduced with a Cre (C) lentivirus 4 days later, and results (R) analyzed 8 days post-electroporation. **j,** mCherry splice junction PCR assay. Black arrows above the mCherry cassette (left) indicate oligo positions. 994bp product: unspliced, 92bp product: spliced. Gel lanes (left to right): molecular weight (mw) ladder, primary neurons electroporated with cL1_spa_ with or without Cre, non-template control, L1_spa_ plasmid positive control, pmCherry plasmid positive, and cL1_spa_ RT^-^ mutant negative control. Red and yellow arrows indicate expected sizes of the spliced and unspliced DNA products, respectively. **k,** Representative confocal images of mCherry immunostaining in primary neurons upon Cre addition. Image insets show cells in merged (top) and single channels for mCherry, Cre, Tub and Hoe (nuclei). Yellow arrows indicate mCherry^+^ neurons. **l,** as in (k) but showing mCherry and PV colocalisation. **m,** as in (k) except showing mCherry immunostaining without Cre. Scale bars in (b-e and k-m) indicate 10µm.

As an orthogonal approach, we electroporated cultured primary mouse neurons with an L1-mCherry retrotransposition reporter based on L1_spa_, a mobile mouse L1 expressed from its native monomeric 5′UTR promoter^28–30^. Whilst we observed PV^+^/mCherry^+^ cells (**Extended Data Fig. 2a,b**), the vast majority of mCherry^+^ cells died, an outcome we did not observe for an L1_spa_ ORF2p reverse transcriptase (RT) mutant (**Extended Data Fig. 2c**). A Cre-LoxP conditional version of the L1-mCherry reporter, which we called cL1_spa_ (**Fig. 1i**), circumvented this apparent toxicity and retrotransposed in primary neurons (**Fig. 1j**-**m**). Nearly all present PV^+^ cells were also mCherry^+^ (**Fig. 1l**). Retrotransposition did not occur in the absence of Cre (**Fig. 1m**) or using an cL1_spa_ ORF2p RT mutant (**Fig. 1j**). As a complementary approach, we used *in utero* electroporation to deliver to the embryonic hippocampus a codon-optimized synthetic L1_spa_^31^ bearing an EGFP reporter (**Extended Data Fig. 3a**). We observed occasional hippocampal EGFP^+^/PV^+^ neurons in neonates (**Extended Data Fig. 3b**). No EGFP^+^ cells were present when the reporter, with disabled ORF2p endonuclease and RT activities^31^, was electroporated into the contralateral hemisphere (**Extended Data Fig. 3c**). Disparate mouse and human L1 reporters delivered by distinct methods thus retrotransposed in the PV^+^ neuron lineage *in vivo* and *in vitro*.

We next measured endogenous L1 expression in PV^+^ neurons. Firstly, we designed a custom single-molecule RNA fluorescence *in situ* hybridization (FISH) probe against the monomeric 5ʹUTR of the mouse L1 T_F_ subfamily^19,28,32^ (**Extended Data Fig. 4a**). With multiplexed RNA FISH, we counted cytoplasmic L1 and PV mRNA puncta in adult β-tubulin (Tub) immunostained neurons (**Fig. 2a** and **Extended Data Fig. 4c,d**). L1 T_F_ transcription was significantly higher in PV^+^ neurons, compared to PV^-^ neurons, in CA (*P*=0.01) (**Fig. 2b**) and DG (*P*=0.009) (**Fig. 2c**). A second L1 T_F_ 5ʹUTR RNA FISH probe (**Extended Data Fig. 4b**) also showed L1 mRNA enrichment in hippocampal and cortical PV^+^ neurons (**Extended Data Fig. 5**). Control experiments conducted in mouse and human cells confirmed the L1 T_F_ RNA FISH probes did not detect DNA (**Extended Data Fig. 4e**) and were specific to L1 T_F_ mRNA (**Extended Data Fig. 4f,g**). Secondly, we used TaqMan qPCR to measure L1 expression in hippocampal PV^+^ neurons and PV^-^ populations (PV^-^, PV^-^/Tub^+^, PV^-^/Tub^-^) sorted from pooled neonate littermates (**Extended Data Fig. 6**). Three qPCR primer/probe combinations (**Extended Data Fig. 4h**) detecting the L1 T_F_ 5ʹUTR each indicated higher expression in PV^+^ neurons than in PV^-^ cells (**Fig. 2d** and **Extended Data Fig. 7a,b**). By contrast, qPCR targeting the L1 T_F_ ORF2 region, expected to mainly detect immobile 5ʹ truncated L1s incorporated in other cellular mRNAs, showed no difference between PV^+^ and PV^-^ cells (**Extended Data Fig. 7c**). Thirdly, 5′RACE upon bulk adult hippocampus or sorted neonate PV^+^ neurons indicated the predominant transcription start sites of full-length L1 mRNAs initiated within L1 T_F_ family 5ʹUTR sequences (**Fig. 2e**). Fourthly, immunostaining using an antibody specific to mouse L1 ORF1p (**Extended Data Fig. 8**), indicated ORF1p expression (**Fig. 2f**) was significantly (*P*=0.0001) higher in PV^+^ neurons than PV^-^ neurons, in CA (**Fig. 2g**) and DG (**Fig. 2h**). Finally, given environmental stimuli can alter neural circuits involving PV^+^ neurons^27^, we analyzed L1 activity using L1 ORF1p immunostaining and L1 T_F_ RNA FISH in adult mice housed in standard (STD), voluntary exercise (EXE) and enriched (ENR) environments, and observed no consistent differences amongst these experimental groups (**Extended Data Fig. 9**, **Extended Data Fig. 10**). We concluded PV^+^ neuron enrichment for L1 T_F_ mRNA and protein expression was robust yet not significantly impacted by exercise or environmental enrichment.

**Fig. 2:**
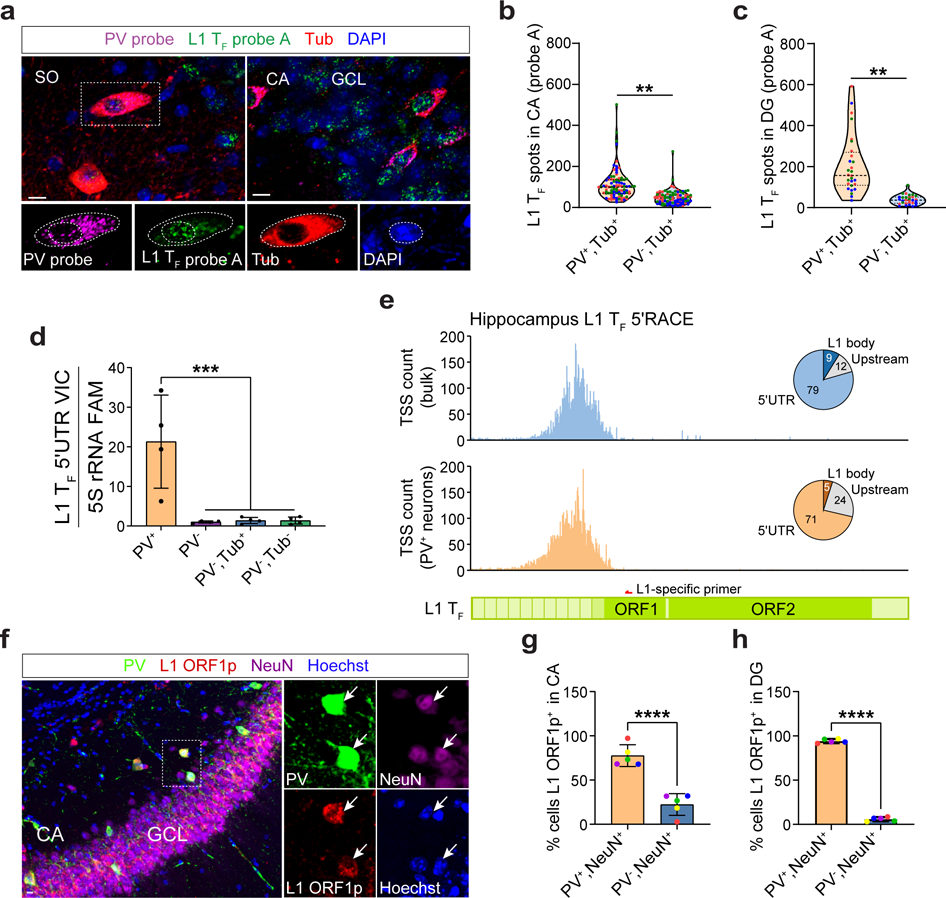
L1 mRNA and ORF1p are abundant in PV^+^ neurons. **a**, Representative maximum intensity projection (MIP) confocal image of a coronal hippocampus section showing L1 T_F_ (green) and PV (magenta) transcripts detected by RNA FISH, β-tubulin (Tub, red) immunohistochemistry and DAPI staining (blue). Image insets show higher magnification of a selected PV^+^ neuron (dashed rectangle). Dashed lines in image insets demark nuclear and cellular boundaries defined for PV and L1 mRNA quantification. Scale bar: 10μm. CA: cornu ammonis, GCL: granular cell layer, SO: stratum oriens. **b,** L1 T_F_ RNA FISH spot (puncta) count per cell in CA PV^+^/Tub^+^ and PV^-^/Tub^+^ neurons. \*\**P*=0.01, PV^+^/Tub^+^ *n*(cells per mouse)=30, PV^-^/Tub^+^ *n*=30, *N*(mice)=3. Cells from different mice are color coded. **c,** As for (b), except for DG. \*\**P*=0.009, PV^+^/Tub^+^ *n*=10, PV^-^/Tub^+^ *n*=10, *N*=3. **d,** Multiplexed TaqMan qPCR measuring mRNA abundance of the L1 T_F_ monomeric 5ʹUTR (VIC channel) relative to 5S rRNA (FAM channel) in PV^+^, PV^-^, PV^-^/Tub^+^ and PV^-^/Tub^-^ cell populations. Cells were sorted from pooled neonate (P0) litter hippocampi. \*\*\**P*=0.001, *N*=4 litters (one-way ANOVA with Tukey’s post-hoc test). Data are represented as mean ± SD. **e,** Transcription start site (TSS) usage within full-length L1 T_F_ copies, detected by L1-specific 5′RACE. RNA was obtained from bulk (blue, top) and sorted PV^+^ hippocampal cells (orange, bottom). Pie charts indicate the percentages of L1 T_F_ mRNAs initiating at upstream TSSs, or TSSs in the L1 T_F_ 5′UTR or body (ORF1). The L1 T_F_ diagram provided underneath indicates the position of the L1-specific primer used for 5′RACE. **f,** Endogenous ORF1p expression in PV^+^ neurons. MIP confocal image of a hippocampus coronal section showing PV (green), ORF1p (red) and NeuN (magenta) colocalization. Insets show a higher magnification view of a selected PV^+^/ORF1p^+^/NeuN^+^ neuron. Hoechst stains nuclei. Scale bar: 10μm. **g,** Percentages of PV^+^/NeuN^+^ and PV^-^/NeuN^+^ neurons expressing ORF1p in hippocampus CA (regions 1-3). \*\*\*\**P*=0.0001, *n*=1,017 average cells per mouse, *N*=5. **h,** As for (g), except in DG. \*\*\*\**P*=0.0001, *n*=719, *N*=5. Note: Significance testing in (b), (c), (g) and (h) was via two-tailed t test, comparing animal or litter mean values.

YY1 and members of the SOX transcription factor family regulate L1 promoters^6,22,23,32–35^. SOX2, for example, represses the youngest human-specific L1 (L1Hs) subfamily in pluripotent and neural stem cells by binding two SOX motifs (first site: +470 to +477, second site: +570 to +577) located in the L1Hs 5ʹUTR^22,23,34^ (**Fig. 3a**). SOX2 downregulation during neurodifferentiation is proposed to facilitate L1 retrotransposition^22,23^. It is unknown whether other SOX proteins then replace SOX2 in binding the L1Hs 5ʹUTR. Crucially, SOX6 coordinates a major transcriptional program of the embryonic and adult brain downstream of LHX6, is necessary for PV^+^ neuron development^8–10^, can bind the first SOX site (**Extended Data Fig. 11a**)^36^ and, of the major hippocampal neuron types, is the only SOX protein specific to PV^+^ neurons (**Extended Data Fig. 11b**). cL1_spa_ reporter assays conducted in primary mouse neurons (**Extended Data Fig. 2d**), analyses of human and mouse ATAC-seq datasets^37,38^ (**Extended Data Fig. 11a,c**), LHX6 overexpression and knockout experiments^39,40^ (**Extended Data Fig. 11d,e**) and SOX6 overexpression experiments performed in cultured mouse primary neurons (**Extended Data Fig. 12**) each gave results congruent with SOX6 activation of both L1 expression and the PV^+^ neuron transcriptional program.

**Fig. 3:**
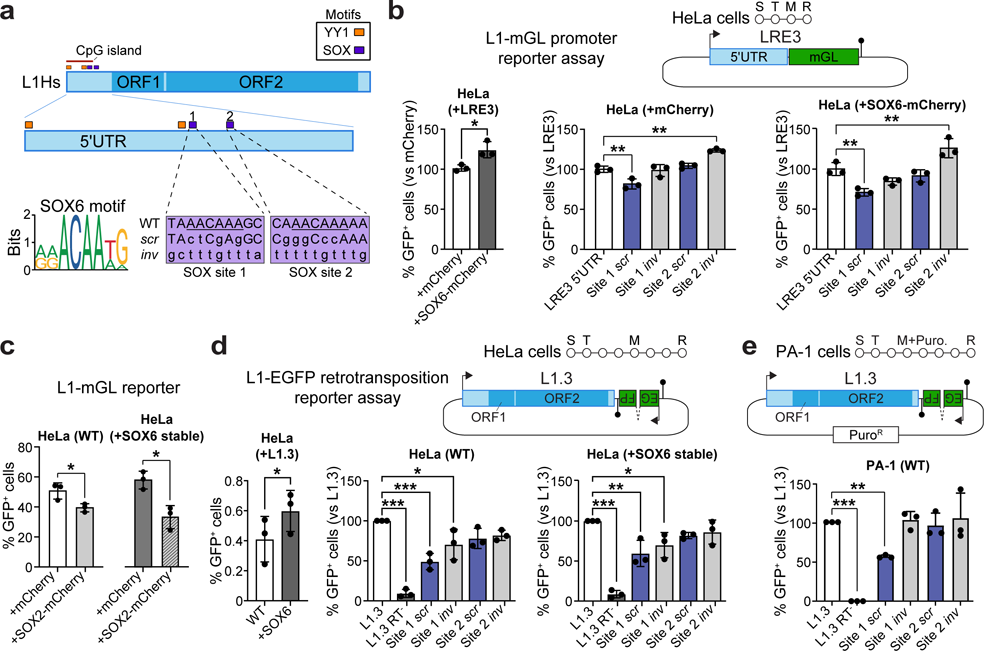
SOX motif-dependent L1 activation by SOX6. **a**, L1Hs schematic. 5ʹUTR embedded YY1- (orange) and SOX-binding (purple) sites are shown, with the latter numbered 1 and 2 and corresponding to L1Hs positions +470 to +477 and +570 to +577, respectively. These SOX motifs were scrambled (*scr*) or inverted (*inv*) in our L1 reporter assays. Site 1 more closely matched the JASPAR^104^ SOX6 binding site motif. **b,** L1 promoter assay. The native 5ʹUTR of the highly mobile L1Hs element LRE3 (top) was used to promote mGreenLantern (mGL) expression (S, seeding; T, transfection; M, change of media; R, result analysis; filled lollipop, polyadenylation signal). Promoter strength is measured as the percentage of GFP^+^ sorted cells. LRE3 5ʹUTR plasmids, including those with scrambled or inverted SOX motifs, were co-transfected into HeLa cells with mCherry (middle) or SOX6-mCherry (right) expression vectors, with a higher percentage of GFP^+^ cells observed in the latter experiment (left). **c,** The reporter from (b) was co-transfected with mCherry or SOX2-mCherry expression vectors into wild-type (WT) and SOX6 stably-overexpressing HeLa cells. **d,** L1 retrotransposition assay. The assay design (top) shows a highly mobile L1Hs element, L1.3 expressed from its native promoter (black arrow) and tagged with an enhanced green fluorescent protein (EGFP) cassette activated upon retrotransposition and driven by a CBh promoter (S, seeding; T, transfection; M, change of media; R, result analysis; filled lollipop, polyadenylation signal. Retrotransposition efficiency is measured as the percentage of GFP^+^ sorted cells. Plasmids included positive (L1.3) and negative (L1.3 RT^-^, D702A mutant) controls and L1.3 sequences with scrambled or inverted SOX motifs. Each element was assayed in WT (middle) and SOX6 stably-overexpressing HeLa cells (right), with a higher percentage of GFP^+^ cells observed in the latter experiment (left). **e,** As for (d), except conducted in WT PA-1 cells using a CMV promoter-driven EGFP cassette and cells were selected for puromycin resistance (Puro^R^). Note: histogram data in (b-e) are represented as mean ± SD with *n*=3 biological replicates. Significance testing in (b), (d) and (e) was via one-way ANOVA against the corresponding positive control (LRE3 5′UTR or L1.3) with Dunnett’s multiple comparison test (middle, right, or only panel) or paired two-tailed t test (left panel). Significance testing in (c) was via paired two-tailed t test. \**P*<0.05, \*\**P*<0.01, \*\*\**P*<0.001.

To dissect L1 activation by SOX6, we generated an L1-mGreenLantern (L1-mGL) promoter reporter based on the 5′UTR of LRE3, another highly mobile human L1^41^. When this L1-mGL reporter was co-transfected into cultured HeLa cells with a SOX6-mCherry expression plasmid, 22% more GFP^+^ cells were observed on average than when cells were co-transfected with an mCherry control plasmid (**Fig. 3b**), a significant difference (*P*=0.02). Inverting either LRE3 5ʹUTR SOX site, or scrambling of the second site, did not reduce L1-mGL activity (**Fig. 3a**,**b**). Scrambling the first SOX site however reduced the percentage of GFP^+^ cells by 18% (*P*=0.004), without SOX6 overexpression, and this reduction was greater (29%, *P*=0.004) when cells were co-transfected with the SOX6-mCherry plasmid (**Fig. 3b**). In agreement with prior reports^22,23^, co-transfection with a SOX2 expression plasmid fully negated SOX6 activation of the L1-mGL reporter (**Fig. 3c**). Next, employing an L1-EGFP retrotransposition reporter^14,24,42^ to assay L1.3^25,26^ mobility in cultured HeLa cells, we found stable SOX6 overexpression significantly (46%, *P*=0.049) increased the percentage of GFP^+^ cells. Scrambling the first SOX site consistently reduced L1.3 retrotransposition efficiency by ∼50%, in HeLa cells with and without stable SOX6 overexpression (**Fig. 3d**) and in cultured PA-1 embryonal carcinoma cells (**Fig. 3e**). These results demonstrated SOX6 activation of L1 promoter and retrotransposition reporters, dependent on the first L1Hs 5ʹUTR SOX site and attenuated by SOX2 expression.

Despite apparent potential for SOX6-mediated L1 transcriptional activation, DNA methylation in somatic cells is expected to silence L1 promoters^6,43,44^. Therefore, to further probe the apparent specificity of L1 transcription to PV^+^ neurons, we performed L1 T_F_ 5ʹUTR monomer bisulfite sequencing^30,35,45^ on neonate hippocampal cell populations. L1 T_F_ was significantly (*P*=0.03) less methylated on average in PV^+^ neurons (83.9%) than in PV^-^ neurons (91.8%) (**Fig. 4a,b**). Unmethylated L1 T_F_ monomers were only observed in PV^+^ neurons (**Fig. 4a,c**). DNMT1, DNMT3A and MeCP2 effect methylation-associated transcriptional repression in PV^+^ neurons^46–49^. These genes all expressed significantly (*P*<0.05) less mRNA in neonate PV^+^ neurons than in PV^-^ neurons (**Fig. 4d,e** and **Extended Data Fig. 13a**). MeCP2 protein expression was on average 10.5% lower in adult PV^+^ neurons, compared to PV^-^ neurons (*P*=0.0007) (**Extended Data Fig. 13b,c**). L1 repression thus appeared broadly relaxed in PV^+^ neurons.

**Fig. 4:**
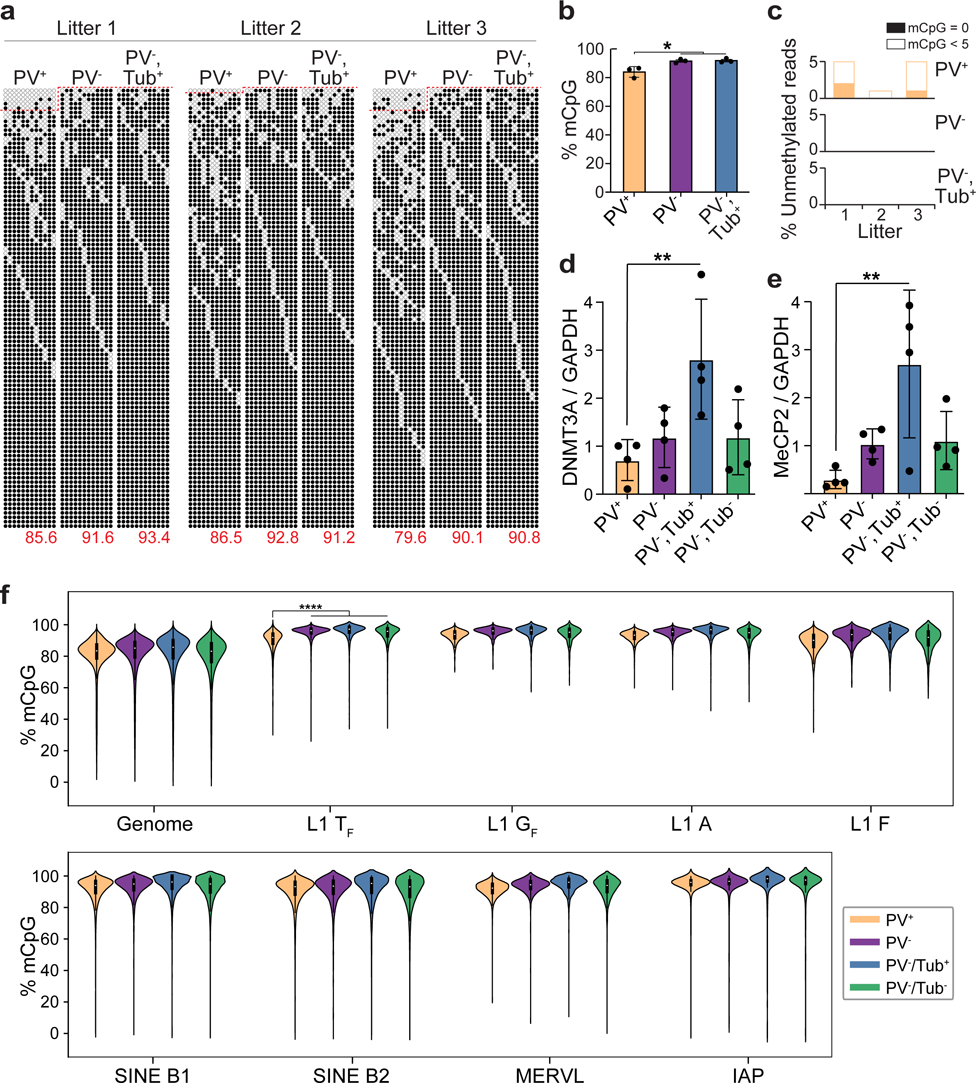
Global L1 T_F_ promoter hypomethylation in PV^+^ neurons. **a**, Targeted bisulfite sequencing of L1 T_F_ promoter monomer CpG islands was performed on PV^+^, PV^-^ and PV^-^/Tub^+^ cells sorted from pooled hippocampal tissue from each of three neonate (P0) litters. Each cartoon displays 100 non-identical randomly selected sequences, where methylated CpGs (mCpGs) and unmethylated CpGs are represented by black and white circles, respectively, as well as the overall mCpG percentage (red numbers). Amplicons above the dotted red line contain <5 mCpGs. **b,** L1 T_F_ monomer methylation was significantly lower (\**P*<0.05, two-tailed t test) in PV^+^ neurons than in either PV^-^ or PV^-^/Tub^+^ cells. **c,** Fully (mCpG=0) and nearly (mCpG<5) unmethylated L1 T_F_ monomers were only found in PV^+^ neurons. **d,** DNMT3A mRNA abundance measured by qPCR in hippocampal cell populations, relative to GAPDH. \*\**P*<0.01, one-way ANOVA with Dunnett’s multiple comparison test only to the PV^+^ population, *N*=4 litters. **e,** As for (d), except for MeCP2. **f,** CpG methylation ascertained by barcoded ONT sequencing upon matched hippocampal PV^+^ and PV^-^ cells from 11 separate neonate litter pools as well as PV^-^/Tub^+^ and PV^-^/Tub^-^ cells from one of those pools. Results are shown for the whole genome (divided into 10kbp windows), and for CpG dinucleotides located within the 5′UTR of T_F_, G_F_, A-type and F-type L1s >6kbp, or within B1 (>140bp) and B2 (>185bp) SINEs, and MERVL MT2 (>470bp) and IAP (>320bp) long terminal repeat sequences. Each included element accrued at least 20 methylation calls and 4 reads in each cell population. \*\*\*\**P*<0.0001, Kruskall-Wallis test comparing the PV^+^ population (*N*=11 litter means) against the three PV^-^ populations combined (*N*=13 litter means) followed by Dunn’s multiple comparison test. Note: histogram data are represented as mean ± SD.

Long-read Oxford Nanopore Technologies (ONT) sequencing allows genome-wide analysis of TE family methylation, as well as that of individual TE loci^30,35,50^. We ONT sequenced PV^+^, PV^-^, PV^-^/Tub^+^ and PV^-^/Tub^-^ cells from neonatal hippocampus samples at ∼16× average genome-wide depth, and ∼29× and ∼34× respectively for the PV^+^ and combined PV^-^ populations (**Supplementary Table 1**). Genome wide, DNA methylation was consistently, if subtly, lower in PV^+^ neurons than in PV^-^ cells (**Fig. 4f**). Amongst the potentially mobile TEs surveyed, only the L1 T_F_ subfamily was significantly (*P*<0.0001) less methylated in PV^+^ cells than in the PV^-^ populations (**Fig. 4f**). L1 loci supplied the vast majority (80%) of differentially methylated TEs (**Fig. 5a**). Comparing PV^+^ and PV^-^ cells, all 638 differentially methylated (*P*<0.01) full-length L1 T_F_ loci were also less methylated in the former population (**Supplementary Table 2**). Those genes containing at least one such L1 were significantly enriched (>20-fold, *P*<0.01) for neurodifferentiation, regulation of GABAergic synaptic signaling, and cell-cell adhesion gene ontologies^51^.

**Fig. 5:**
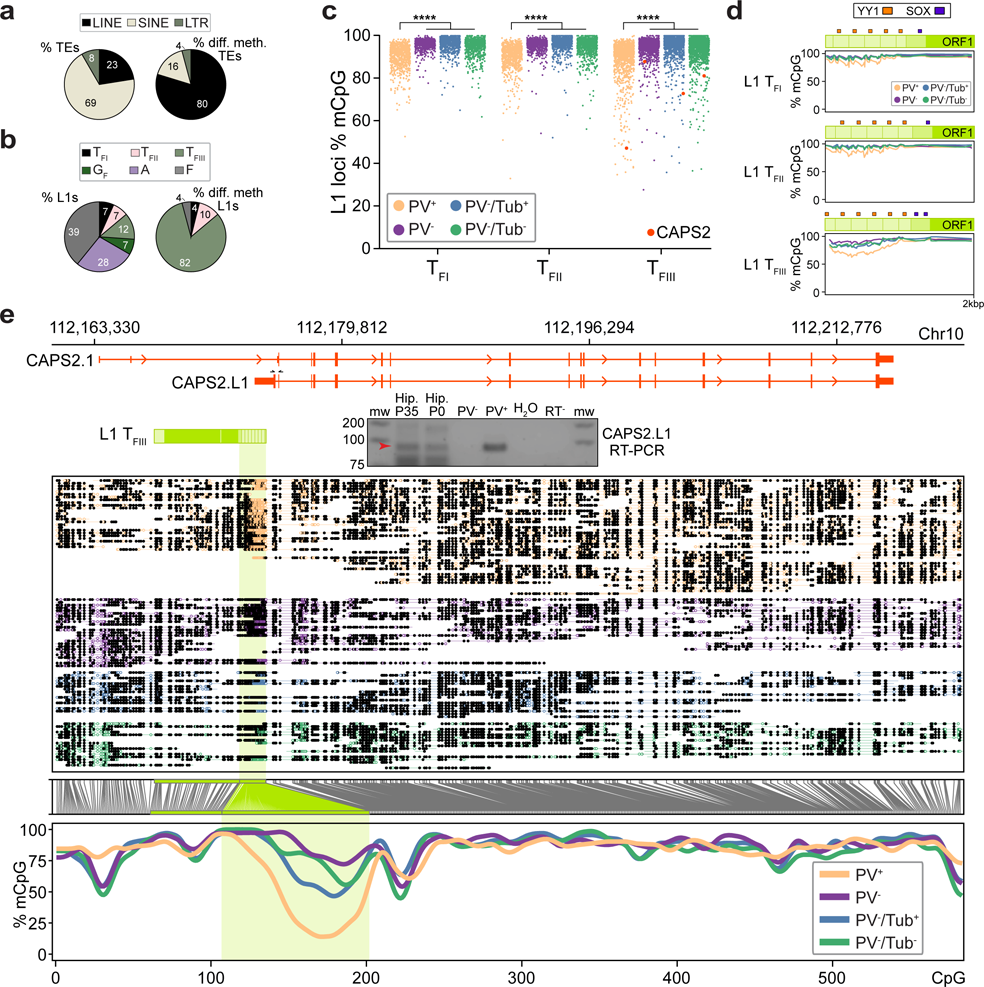
PV^+^ neuron genes harbor hypomethylated L1 T_F_ promoters. **a**, Composition of all young (left) and differentially methylated (*P*<0.01) young (right) TEs, by superfamily. Note: MERVL and IAP are LTR retrotransposons. **b,** As per (a), except showing the breakdown of young L1 subfamilies (left) and their contribution to the 50 differentially methylated loci showing the largest absolute change in methylation percentage. **c,** L1 T_FI_, T_FII_ and T_FIII_ subfamily CpG methylation strip plots for PV^+^, PV^-^, PV^-^/Tub^+^ and PV^-^/Tub^-^ cells, as represented collectively by the L1 T_F_ violin plot in Fig. 4f. Each point represents an L1 locus, with an example intronic to CAPS2 highlighted by an orange dot. \*\*\*\**P*<0.0001, Kruskall-Wallis test on PV^+^ population (*N*=11 litter means) versus three PV^-^ populations combined (*N*=13 litter means) followed by Dunn’s multiple comparison test. **d,** Composite DNA methylation profiles for the L1 T_F_ subfamilies displayed in (c), representing the mean methylation for CpGs within the first 2kbp of elements with 6 monomers. Darker green shading represents the non-monomeric 5′UTR region. **e,** Methylation profile of the CAPS2 locus obtained from ONT sequencing. The first panel shows a full-length L1 T_FIII_ with intact ORFs, as highlighted in (c), orientated antisense to the first intron of the canonical CAPS2.1 isoform. ENCODE long-read transcriptome sequencing of hippocampus tissue (ENCLB505CBY) indicated a chimeric transcript, labeled here CAPS2.L1, spliced into CAPS2.1 exon 3 and encoding an ORF in frame with the CAPS2.1 ORF. The gel image displays PCR products generated using primers specific to CAPS2.L1 (marked by opposing black arrows), with input template cDNA from bulk adult (P35) hippocampus (5ʹRACE) and neonate (P0) hippocampus (reverse transcribed total RNA from bulk and sorted PV^+^ and PV^-^ cells). A red arrow indicates on-target products confirmed by capillary sequencing. The second panel displays aligned ONT reads, with unmethylated CpGs colored in orange (PV^+^), purple (PV^-^), blue (PV^-^/Tub^+^) and green (PV^-^/Tub^-^), and methylated CpGs colored black. The third panel indicates the relationship between CpG positions in genome space and CpG space, including those corresponding to the L1 T_FIII_ 5ʹUTR (shaded light green). The fourth panel indicates the fraction of methylated CpGs for each cell type across CpG space.

Notably, the T_F_ subfamily can be further divided into three subgroups, denoted T_FI_, T_FII_, and T_FIII_ (**Fig. 5d**), where T_FIII_ is the oldest and diverges in its 5ʹUTR when compared to T ^52^. We found by far the highest fraction (82%) of strongly demethylated L1s corresponded to the T_FIII_ subfamily (**Fig. 5b**-**d**). Strikingly, we identified significantly (*P*<0.01) hypomethylated L1 T_FIII_ copies with intact ORFs in the introns of genes expressed in PV^+^ neurons, such as CAPS2 (calcyphosphine 2)^53^, CHL1^54^, ERBB4^55^ and NPSR1^56^ (**Fig. 5e**, **Extended Data Fig. 11b**, **Extended Data Fig. 14** and **Supplementary Table 2**). In CAPS2, which expresses an EF-hand calcium-binding protein in the same family as parvalbumin^53^, the L1 5ʹUTR was completely unmethylated in numerous PV^+^ neurons (**Fig. 5e**). Analysis of ENCODE PacBio long-read hippocampus transcriptome sequencing^57^ revealed a transcript initiated within and antisense to the L1 5ʹUTR and spliced downstream into a CAPS2 exon on the same strand. We termed this chimeric transcript CAPS2.L1. By 5ʹRACE and RT-PCR, we reliably detected CAPS2.L1 in adult and neonate hippocampus tissue, and in PV^+^ cells, but not PV^-^ cells (**Fig. 5e**). The predicted ORF for CAPS2.L1 was in frame with the canonical CAPS2 ORF and incorporated a novel N-terminal sequence (**Fig. 5e**). When introduced as an expression vector (**Fig. 6a**) into mouse N2a neuroblastoma cells, a tractable model of neurodifferentiation, CAPS2.L1 significantly increased neuron branching and neurotrophin-3 (NTF3) release, compared to CAPS2.1 and controls (**Fig. 6a**-**c**). These results suggested CAPS2.L1 enhanced neuron morphological complexity and function. As the annotated CAPS2 promoter was fully methylated in PV^+^ neurons (**Fig. 5e**), we concluded CAPS2.L1 is likely the canonical CAPS2 transcript isoform in mouse PV^+^ neurons, previously overlooked due to its initiation in the L1 T_FIII_ sequence. Finally, including the CAPS2 example, we identified 43 young mouse L1s whose 5ʹUTR promoted expression of a spliced transcript annotated by GenBank or detected by the abovementioned ENCODE PacBio datasets (**Supplementary Table 3**). These results suggested unmethylated L1s can promote transcription of genes required for mouse PV^+^ neuron development and function.

**Fig. 6:**
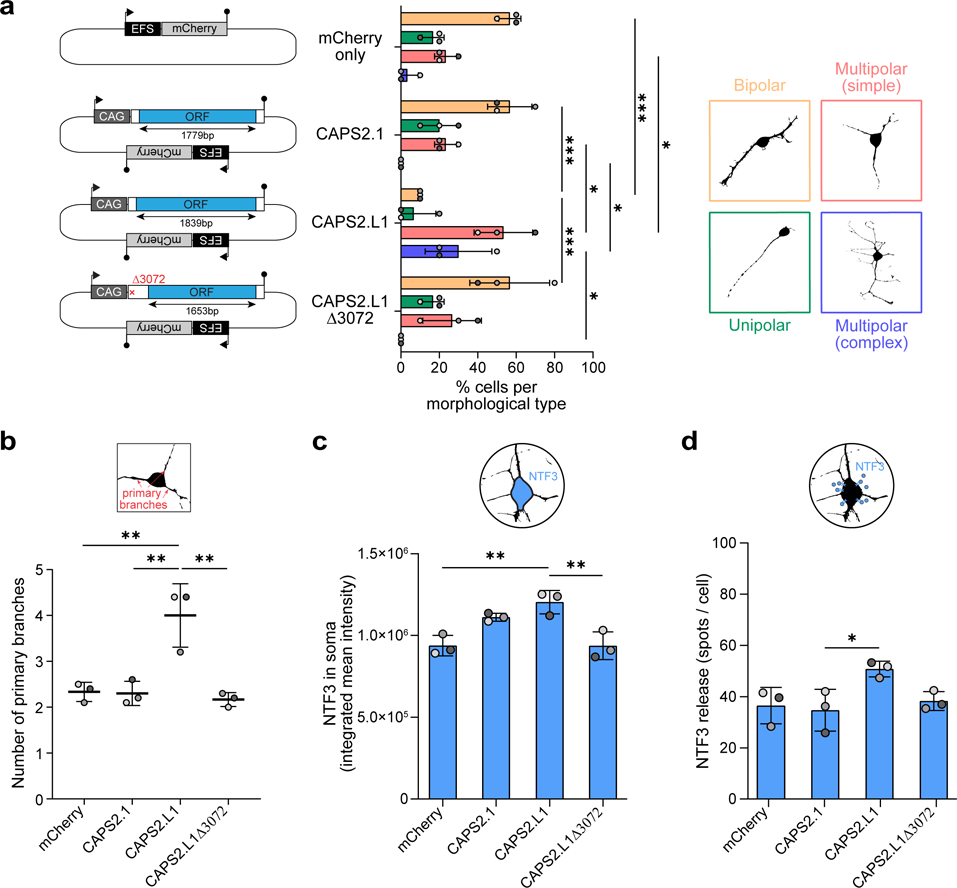
CAPS2.L1 enhances neuron morphological complexity and NTF3 release. **a**, CAPS2 mRNA expression construct design (left). Each is expressed from a CAG promoter. The CAPS2.L1 ORF (blue) is larger than that of CAPS2.1 and encodes an alternative N-terminus, whereas the CAPS2.L1Δ3072 construct is identical to CAPS2.L1 except with a 4bp deletion at position 3072 that results in a truncated CAPS2.L1 ORF. All constructs have an mCherry cassette driven by an EFS promoter on their backbone. An empty mCherry vector was used as a transfection control. Promoters are represented by black arrows. Filled lollipops indicate polyadenylation signals. N2a neuroblastoma cells were transfected with each construct (left) and differentiated, and the morphology type (right) of mCherry^+^ cells quantified (middle). Observed morphological types were classified as: (i) bipolar (orange), (ii) unipolar (green), (iii) multipolar (simple) (red), or (iv) multipolar (complex) (blue). Each histogram data point represents the average of values from an independent experiment. *N*=3 experiments, *n*=10 cells per experiment. **b,** CAPS2 constructs assayed as in (a) but here showing the number of primary branches per cell. **c,** NTF3 integrated mean immunofluorescence intensity in cell soma. *N*=3 experiments, *n*=95 cells per experiment. **d,** NTF3 release quantified as NTF3 spots per cell, outside of the soma. Analysis was performed on high magnification confocal images within a fixed radius set around the cell soma. *N*=3 experiments, *n*=8-10 cells per experiment Note: in each histogram, experimental replicates are colored different shades of gray and data are represented as mean ± SD. Significance values were calculated based on the averages of independent experiments via one-way ANOVA with Tukey’s post-hoc test. \**P*<0.05, \*\**P*<0.01, \*\*\**P*<0.001.

In sum, this study reveals L1 activity in the PV^+^ neuron lineage governed by SOX6 (**Extended Data Fig. 15**). PV^+^ neurons are “node” cells that connect neural circuits associated with memory consolidation and other core cognitive processes^58,59^. The potential for L1 mobility as a consequence of PV^+^ neuron genes incorporating unmethylated retrotransposition-competent L1s is notable given the proposed roles for stochastic L1-mediated genome mosaicism in the brain^22,60–62^. Our results do not however preclude other neuronal subtypes or brain regions from expressing L1 mRNAs and proteins, or supporting L1 retrotransposition. Engineered L1 reporter experiments have thus far generated data congruent with endogenous L1 mobility in the early embryo^18,19,21^, neurons^4–7,22,23,63^ and cancer^14,64,65^. While we and others have mapped endogenous L1 retrotransposition events in human^4–6^ and macaque^7^ neurons, the composition of the L1 T_F_ 3ʹUTR appears to severely impede such analyses in mouse^19,45,66^. A cL1_spa_ mouse model could in the future be used to evaluate mouse L1 mobilization in the PV^+^ neuron lineage *in vivo*.

As highlighted here and elsewhere^1,3,11,13,67–69^, retrotransposons can be integrated into transcriptional programs guiding cell fate. Hypomethylated L1s could also influence the regulatory architecture of adjacent genes by attracting SOX proteins and their cofactors^34^. DNA methyltransferase activity appears to be moderately attenuated in PV^+^ neurons. However, to explain L1 hypomethylation in this context, we favor a model where some L1 loci escape embryonic methylation and are transcribed^6,35^. Histone modifications associated with active transcription, perhaps enhanced by SOX6, then counteract their methylation in PV^+^ neurons^70–72^. These heritable ‘escapee’ L1s, along with their transcriptional, regulatory and mobilization potential, are subject to evolutionary selection. Somatic retrotransposition could hence signify niches of L1 cis-regulatory innovation, as revealed here in the PV^+^ neuron lineage.

## Acknowledgements

We thank Jef D. Boeke, Jose L. Garcia-Perez, Haig H. Kazazian and John V. Moran for sharing L1SM, LRE3, L1_spa_ and L1.3 plasmids. We thank Wenfeng An, Pankaj Sah, Ryan Lister, Robert Sullivan, Sam van de Wakker, Arnaud Gaudin, Seth W. Cheetham, and members of the Faulkner laboratory for helpful discussions. We acknowledge the QBI and TRI Flow Cytometry suites for technical advice, the Garvan Sequencing Platform and the Australian Genome Research Facility for long-read sequencing, the UQ School of Biomedical Sciences Viral Vector Core for Cre lentivirus synthesis, and the QBI Advanced Microscopy Facility for technical assistance and equipment, supported by ARC LIEF grant LE130100078. This work was supported by NHMRC-ARC Dementia Research Development Fellowship GNT1108258 and DFG fellowship BO4460/1-1 (G.O.B.), Australian Government Research Training Program (RTP) Scholarships (J.M.B. and M.E.F.), NHMRC Investigator Grants GNT1176574 (N.J.), GNT1173476 (S.R.R.) and GNT1173711 (G.J.F.), ARC Discovery Project DP200102919 (S.R.R. and G.J.F.), NHMRC Project Grants GNT1106206 (G.J.F., A.J.H., L.M.P.) and GNT1126393 (G.J.F.), a CSL Centenary Fellowship (G.J.F), Andalusian Government EMERGIA20_00225 and Spanish National Research Agency RYC2021_031920-I grants (F.J.S-L), and the Mater Foundation.

## Author contributions

G.O.B. and G.J.F. designed the project and wrote the manuscript. G.O.B., J.M.B., M.E.F., F.J.S-L., J.d.l.R.B., J.R., M.A.R., L.R.F., C.G., P.G., L-G.B, P.A., D.J.D.C.G., L.C., C.C.B., V.B., S.M., M-J.H.C.K., C.J.L. and K.S. performed experiments. G.O.B., J.M.B., M.E.F., N.J., P.K., A.D.E., D.J.J., S.R.R., A.J.H. and G.J.F. analyzed the data. G.O.B., L.M.P., S.R.R., A.J.H. and G.J.F. provided resources.

## Data availability

ONT sequencing data (.fastq and .fast5) generated from sorted hippocampal cell populations are available from the European Nucleotide Archive (ENA) under accession number PRJEB47835.

## Materials availability

L1.3 retrotransposition reporter and LRE3 promoter reporter constructs carrying mutant SOX6 binding sites are available from Geoffrey J. Faulkner and require a material transfer agreement.

### Code availability

Nanopore methylation analyses were performed with MethylArtist (https://github.com/adamewing/methylartist). Bisulfite sequencing results were visualized with QUMA (http://quma.cdb.riken.jp/). RNA-seq and ATAC-seq datasets were analyzed by pipelines joining together in serial published bioinformatic tools (see **Methods**). A custom Python script (https://gist.github.com/adamewing/331e1780975969afcebc2996ddb8c7a2) was written to analyze scATAC-seq profiles of the L1Hs promoter.

## Methods

### Cultured cell L1 retrotransposition reporter assay

L1 retrotransposition efficiency was measured via an enhanced green fluorescent protein (EGFP) L1 reporter system in cultured HeLa and PA-1 cells, as described previously^6,24,42,73^. These assays employed the pCEP4_L1_eGFPI plasmid^6^ and tested the mobility of wild-type and reverse transcriptase mutant^14^ (D702A) L1.3^25,26^ sequences expressed from the native L1 promoter, as well as wild-type L1.3 sequences with their 5′UTR SOX binding sites scrambled or inverted^34^. Each SOX site mutation was introduced by fusion PCR to build an L1 fragment flanked by NotI (5′) and AgeI (3′) sites from two amplicons with overlapping primers that included the desired mutation. The complete amplicon was cloned within these sites in the original backbone and its sequence was verified by capillary sequencing. In this L1 reporter^6^, the entire L1 3ʹUTR, with the thymine base deleted from within its native polyadenylation signal, preceded an EGFP reporter cassette activated only upon retrotransposition. EGFP expression was driven by a CBh (in HeLa) or CMV promoter (in PA-1). GFP^+^ cells were counted via flow cytometry. The L1 plasmid backbone incorporated a puromycin resistance gene. Three biological replicate assays were performed, each consisting of 3 wells per condition (technical replicates), on different days.

HeLa-JVM cells^14^ (obtained from the laboratory of John V. Moran) were cultured at 37°C and 5% CO_2_ in HeLa complete medium (DMEM, Life Technologies, Cat# 11960044) supplemented with 10% FBS (Life Technologies, Cat# 10099141), 1% Glutamax (Life Technologies, Cat# 35050061) and 1% penicillin-streptomycin (Life Technologies, Cat# 15140122). Cells were passaged at 70-80% confluency using 0.25% Trypsin-EDTA (Life Technologies, Cat# 25200072). Briefly, 5×10^4^ HeLa cells were seeded per well of a 6-well plate. Eighteen hours later, cells were transfected with 1µg L1 plasmid per well using 3µL FuGENE HD transfection reagent (Promega, Cat# E2311) and 97µL Opti-MEM (Life Technologies, Cat# 31985047) per well according to the manufacturer’s protocol. Twenty-four hours post-transfection, medium was replaced with HeLa complete medium. No puromycin selection was performed. Medium was replaced every other day, and cells were collected 6 days post-transfection by trypsinization, resuspended in sterile PBS, and analyzed on a BD FACSymphony™ A5 SE Cell Analyzer (BD Biosciences) using FlowJo (version 10.8.1) to determine the percentage of GFP^+^ cells. Untransfected HeLa cells were used to set the GFP^-^ signal level in flow cytometry.

To assess L1 retrotransposition coincident with stable SOX6 overexpression, human SOX6 cDNA (NM_001145819.2) driven by a CBh promoter and mCherry driven by an EFS promoter were inserted using BsrGI and Acc65I restriction enzymes (NEB, Cat# R3575S and R0599S, respectively) into the XLone-GFP plasmid (Addgene, Cat# 96930). HeLa cells were co-transfected with this SOX6 expression plasmid and a hyperactive Piggybac transposase^74^ (HyPBase, a kind gift from the laboratory of Jose M. Polo) at a 3:1 ratio. The Piggybac plasmid harbored a blasticidin resistance gene, allowing transfected cells to be selected using 20µg/mL of blasticidin for 2 weeks post-transfection to establish a stable SOX6 overexpressing HeLa cell line.

PA-1 cells were purchased from the American Type Culture Collection (ATCC), cultured at 37°C and 5% CO_2_, and maintained in Minimum Essential Medium (MEM) with GlutaMAX supplement (Life Technologies), 10% heat-inactivated FBS, 1× non-essential amino acids (NEAA, Life Technologies) and 100U/mL Pen-Strep solution (Life Technologies). 2×10^5^ PA-1 cells were seeded per well of a 6-well plate and transfected as per the HeLa cell experiments above. Medium was replaced daily with PA-1 complete medium supplemented with 0.5µg/mL puromycin the day after transfection, 1µg/mL puromycin each day afterwards, and 500nM trichostatin A (Sigma, Cat# T8552) 12 hours prior to flow cytometry on day 6 after transfection. Flow cytometry employed a BD Accuri C6 Flow Cytometer (BD Biosciences). No untransfected PA-1 cells survived treatment with puromycin, ensuring untransfected cells did not contribute to GFP^-^ cells on the day of analysis. Untransfected PA-1 cells not treated with puromycin were used to set the GFP^-^ signal level in flow cytometry. As a quality check, plasmid transfection efficiencies were calculated by co-transfecting with pCEP-EGFP^75,76^.

### cL1_spa_ retrotransposition in primary neuronal cultures

We designed a Cre-LoxP^77–79^ conditional L1 retrotransposition reporter, cL1_spa_, based on a variant of the mobile L1 T_F_ element L1_spa_^28^. Here, L1_spa_ was contained in a pCEP4 backbone, with ORF1p corrected at two amino acid positions to match the L1 T_F_ subfamily consensus sequence, and had an mCherry indicator cassette driven by an EF-1ɑ promoter embedded in its 3ʹUTR^29,30^. The G-rich region of the 3’UTR (nucleotides 7260-7416) was moved downstream of the reporter cassette, cloned between two BamHI sites before the SV40 polyadenylation signal^29^. To prohibit full-length L1_spa_ mRNA production in the absence of Cre recombinase, between the L1_spa_ 5′UTR promoter and ORF1 we inserted a LSL cassette containing two LoxP sequences (ATAACTTCGTATAGCATACATTATACGAAGTTAT) flanking a 786bp sequence containing three SV40 polyadenylation signals and followed by a Kozak consensus sequence (Vector Builder). We synthesized a fragment containing the 5′UTR, LoxP-Stop-LoxP and ORF1 (GenScript) sequences, flanked by Not1 and PacI restriction sites, and cloned it into the L1_spa_ backbone. We used this construct to electroporate primary neuronal cells using a Neon NxT Electroporation System (ThermoFisher Scientific) and the Neon Transfection System 10µL Kit (ThermoFisher Scientific, Cat# MPK1025,) at 1500V, 10ms, and 3 pulses. 2.5µg of plasmid was used per electroporation, with two electroporations plated per 12-well plate well.

Primary neuronal cultures were prepared from embryonic day (E18) mice by dissecting the cortices and placing them in a papain solution (Worthington Biochemical, Cat# LS003126) dissolved in a basic culture medium (Neurobasal medium (Gibco, Cat# 21103-049), GlutaMAX (Gibco, Cat# 35050061), and Penicillin-Streptomycin (5000 U/mL, Gibco, Cat# 15070063). Cortices were incubated for 20min at 37°C with pipette mixing every 5min. Cells were then filtered through a 70µm cell strainer into an inactivation solution containing albumin-ovomucoid inhibitor (Worthington, Cat#LK003182), DNAseI (1mg/mL, Sigma, Cat# DN-25) and basic culture medium with 5% FBS. Cells were centrifuged at 400g for 2min at room temperature and then resuspended in 5% FBS culture medium and counted. 250,000 cells were washed with PBS twice and used for one electroporation. Cells were then plated on Poly-D-Lysine (1mg/mL, Sigma P64407) coated coverslips. Four days post-electroporation, cells were transduced with a lentiviral construct expressing Cre-T2A-TagBFP driven by an EFS moderate EF-1α core promoter (Vector Builder). Four days post-transduction, cells were processed either for immunostaining or DNA extraction. For immunostaining, culture medium was removed, and cells were immediately fixed in 4% PFA (20min, 4°C) and then washed with PBS. Cells were permeabilized for 5min with PBT (PBS with 0.5% Triton) and then incubated with blocking buffer (10% normal donkey serum (Jackson ImmunoResearch, Cat# AB2337258) in PBT) for 1 hour at room temperature. Cells were incubated overnight with the following primary antibodies: goat anti-tdTomato (SICGEN, Cat# AB8181, 1:2000), mouse anti-β-tubulin (Sigma, Cat# T4026, 1:1000) and either rabbit anti-Cre (Cell Signaling, Cat# 15036, 1:500) or rabbit anti-parvalbumin (Swant, PV27, 1:500). Cells were washed with PBS for 15min, 3 times at room temperature and then incubated with secondary antibodies: Cy3 donkey anti-goat (Jackson ImmunoResearch, Cat# 715-165-150, 1:500), Alexa Fluor 488 donkey anti-mouse (Jackson ImmunoResearch, Cat# 715-546-150, 1:1000) and Alexa Fluor 647 donkey anti-rabbit (Jackson ImmunoResearch, Cat# 711-606-152, 1:1000). Hoechst 33258 (Sigma, Cat# B2883) was added to the secondary antibody mix to stain nuclei. Cells were again washed with PBS for 15min, 3 times at room temperature, then air dried and mounted using an aqueous fluorescence mounting medium (Agilent Dako, Cat# S302380-2). Cells were visualized and imaged using an Olympus UPLXAPO 20×/0.8 NA air objective on a spinning disk confocal microscope (SpinSR10; Olympus, Japan) built around an Olympus IX3 body and equipped with two ORCA-Fusion BT sCMOS cameras (Hamamatsu Photonics K.K., Japan) and controlled by Olympus cellSens software.

Genomic DNA was phenol-chloroform extracted. Oligonucleotide PCR primers spanning the mCherry cassette exon-exon junction (**Supplementary Table 4**) were purchased from Integrated DNA Technologies. PCR reactions were prepared using the MyTaq DNA polymerase kit (Bioline, Cat# BIO-21111). Reactions contained a 5× MyTaq Buffer, 10pmol of each primer, 1µL DNA input template (20ng) and 0.2µL Taq polymerase in 25μL final volume. Cycling conditions were as follows: 95°C for 2min, followed by 30 cycles of 95°C, 15sec; 58°C, 15sec; 72°C, 50 sec and extension 72°C, 5min. PCR products were separated on 1.5% agarose gel.

### Cultured cell L1 promoter reporter assay

The effects of SOX site mutations on L1 promoter activity were assessed in cultured HeLa-JVM cells using the LRE3^41^ 5ʹUTR to drive expression of a mGreenLantern^80^ (mGL) green fluorescent protein. The native LRE3 5ʹUTR and SOX site mutants were ordered as a synthetic double stranded DNA gene blocks from Integrated DNA Technologies and incorporated EcoRI and BamHI restriction sites at their 5ʹ and 3ʹ ends, respectively. These sites were used to clone the LRE3 5ʹUTR into an expression vector, resulting in an L1 promoter reporter. SOX protein overexpression was evaluated using the L1 promoter reporter in HeLa cells in two experiments. In the first experiment, cells were seeded at 1×10^5^ cells/well in a 6-well plate. 16 hours after seeding, cells were co-transfected with 3µL FuGENE HD, 97µL Opti-MEM and a 1:1 ratio of the L1 promoter reporter and a plasmid encoding the human SOX6 (NM_001145819.2) cDNA, driven by a CBh promoter, and including an mCherry cDNA under the control of an EFS promoter to assess transfection efficiency (Vector Builder). This plasmid, without SOX6, was used as a control. The transfection medium was replaced after 24 hours with DMEM-complete medium and the cells were incubated for 24 hours, after which the percentage of GFP^+^ cells was determined by flow cytometry. In the second experiment, WT and stably SOX6 overexpressing cells were seeded at 1×10^5^ cells/well in a 6-well plate. 16 hours after seeding, both cell lines were cotransfected with 3µL FuGENE HD, 97µL Opti-MEM and a 1:1 ratio of the L1 promoter reporter and a plasmid encoding SOX2 cDNA (NM_003106.4) under a CBh promoter (Vector Builder). This plasmid also included an EFS promoter driving the expression of mCherry to monitor transfection efficiency. The same plasmid without SOX2 was used as a control. 24 hours later, the transfection medium was replaced with DMEM-complete medium and cells incubated for 48 hours, after which the percentage of GFP^+^ cells was analyzed by flow cytometry.

### SOX6 expression assay in primary neuronal cultures

Primary neuronal cultures were transduced at day 5 post plating with 1µL/well of 12-well plate using either adeno-associated virus of serotype 2 (AAV2) encoding the mouse SOX6 (NM_011445.4) cDNA or mCherry control at a concentration of >2×10^11^ genome copies (GC)/mL. SOX6 expression was driven by a CBh promoter, and the construct included an mCherry cassette under the control of an EFS promoter to assess transduction efficiency (Vector Builder). 24h after transduction, RNA was extracted using Trizol reagent (Invitrogen Cat# 15596026) following the manufacturer’s protocol. SYBR Green qPCR was prepared using the Power SYBR Green RNA-to-CT 1 step kit (Applied Biosystems, Cat# 4391112) and specific SOX6 primers (mSOX6_F and mSOX6_R) to assess SOX6 overexpression. Reactions contained a 2× Power SYBR Green RT-PCR Mix, 10pmol of each primer, 1µL RNA input template and 1× reverse transcriptase enzyme mix in a 10μL final volume. Cycling conditions were as follows: 48°C for 30min, 95°C for 10min, followed by 40 cycles of 95°C, 15sec; 60°C, 1min. To assess potential DNA contamination, an L1 T_F_ qPCR using primers L1Md_5UTR_F and L1Md_5UTR_R was performed with and without reverse transcriptase. A three or more cycle difference between experiments run with and without reverse transcriptase, and detection after cycle 30 in the latter, was considered as non-DNA contaminated RNA. A TaqMan assay was also performed using Applied Biosystems custom L1 and 5S rRNA TaqMan MGB probes, as listed in **Supplementary Table 4**. TaqMan qPCR reactions contained: 4× TaqPath 1-Step RT-qPCR multiplex reaction master mix (ThermoFisher, Cat# A28521), 4pmol of each primer, 1pmol probe and 1µL RNA (100-150 ng) input template in a 10uL final volume. Cycling conditions were as follows: 37°C for 2min; 50°C for 15min; 95°C for 2min, followed by 40 cycles of 95°C, 3sec; 60°C, 30sec. TaqMan assays for L1 probe conjugated with VIC were multiplexed with an assay for 5S rRNA control, conjugated to 6FAM fluorophores. Primer/probe sequences and the associated detection channels are listed in **Supplementary Table 4**. 72h after transduction, the culture medium was removed, and cells were fixed in 4% PFA (20min, 4°C) and washed with PBS. Immunohistochemistry was then performed as for mouse primary neuronal cultures. For ORF1p immunostaining analysis, Z-stack images were acquired using the SoRa super resolution disk and 3.2× magnification. Image processing and analysis post-acquisition for the morphology analysis were performed using Fiji for Windows (ImageJ 1.52d). ORF1p intensity analysis in mCherry^+^ cells was performed using Imaris 9.5.1 (Bitplane, Oxford Instruments). Integrated mean intensity was calculated as equal to area times mean intensity value and normalized to the mCherry control for the respective experiment.

### L1-EGFP transgenic mice

To trace retrotransposition of an engineered L1 reporter *in vivo*, we generated a new transgenic L1-EGFP mouse line harboring L1.3, with epitope tags on ORF1p and ORF2p and an EGFP indicator cassette^14,24^ embedded in its 3ʹUTR. To assemble the L1 transgene, we cloned the NotI-BstZ17I fragment from pJM101/L1.3-ORF1-T7-ORF2-3×FLAG (containing T7 gene 10 epitope tag on the C-terminus of ORF1 and a 3×FLAG tag on the C-terminus of ORF2) into p99-GFP-LRE3, yielding p99-GFP-L1.3-ORF1-T7-ORF2-3×FLAG. In p99-GFP-L1.3-ORF1-T7-ORF2-3×FLAG, transgene transcription was driven by the native L1.3 promoter, with an SV40 polyadenylation signal (pA) located downstream of the EGFP retrotransposition indicator cassette. The EGFP cassette was equipped with a cytomegalovirus (CMV) promoter and a herpes simplex virus type 1 (HSV) thymidine kinase (TK) polyadenylation signal, facilitating EGFP expression upon genomic integration via retrotransposition. In preparation for pronuclear injection, EGFP-L1.3-ORF1-T7-ORF2-3×FLAG was released by digestion with Not1 and MluI restriction enzymes, separated from the vector backbone on a 0.7% agarose gel, purified by phenol-chloroform extraction, and eluted in microinjection buffer (7.5mM Tris-HCl, 0.15mM EDTA pH7.4). Transgenic L1-EGFP mice were produced by the Transgenic Animal Service of Queensland (TASQ), University of Queensland, using a standard pronuclear injection protocol. Briefly, zygotes were collected from superovulated C57BL/6 females. The microinjection buffer containing EGFP-L1.3-ORF1-T7-ORF2-3×FLAG was then transferred to the zygote pronuclei. Successfully injected zygotes were transplanted into the oviducts of pseudopregnant females. Primers flanking the EGFP cassette were used to screen potential founders by PCR (**Supplementary Table 4**). Identified founder L1-EGFP animals were bred on a C57BL/6 background. All procedures were followed as approved by the University of Queensland Animal Ethics Committee (TRI/UQ-MRI/381/14/NHMRC/DFG and MRI-UQ/QBI/415/17).

### In utero electroporation

Embryonic *in utero* electroporation was employed to simultaneously deliver control (pmCherry) and experimental (L1) plasmids. Here, pmCherry *w*as a 4.7kb plasmid that expressed mCherry fluorescent protein under the control of a CMV promoter (Addgene, Cat# 632524). L1 plasmids consisted of pUBC-L1SM-UBC-EGFP and pMut2-UBC-L1SM-UBC-EGFP. pUBC-L1SM-UBC-EGFP was a derivative of cep99-GFP-L1SM, which contained a full-length codon-optimized synthetic mouse L1 T_F_ element (L1SM, kindly shared by Jef Boeke, NYU Langone)^31^, where mouse ubiquitin C (UBC) promoters were substituted for the CMV promoters used to drive L1SM and EGFP expression in cep99-GFP-L1SM. pMut2-UBC-L1SM-UBC-EGFP was identical to pUBC-L1SM-UBC-EGFP, apart from two non-synonymous mutations in the L1SM ORF2 sequence known to disable ORF2p reverse transcriptase and endonuclease activities. *In utero* electroporation was performed as described previously^81^, with the day of mating defined as embryonic day 0 (E0). Briefly, time-mated pregnant CD1 mice were anesthetized at E14.5 via an intraperitoneal injection of ketamine/xylazine (120mg/kg ketamine and 10mg/kg xylazine). Embryos were exposed via a laparotomy and 0.5-1.0μL of plasmid DNA combined with 0.0025% Fast Green dye, to aid visualization, was injected into the lateral ventricle of each embryo using a glass-pulled pipette connected to a Picospritzer II (Parker Hannifin). Injections involved either combinations of pUBC-L1SM-UBC-EGFP and pmCherry (1μg/μL each) or pMut2-UBC-L1SM-UBC-EGFP and pmCherry (1μg/μL each). Half of the pups from each litter were co-injected with pUBC-L1SM-UBC-EGFP and pmCherry into the left hemisphere and the other half with pMut2-UBC-L1SM-UBC-EGFP and pmCherry into the right hemisphere. Plasmids were directed into the forebrain by placement of 3mm diameter microelectrodes across the head, which delivered 5 (100ms, 1Hz) approximately 36V square wave pulses via an ECM 830 electroporator (BTX). Once embryos were electroporated, uterine horns were replaced inside the abdominal cavity and the incision sutured closed. Dams received 1mL of Ringer’s solution subcutaneously and an edible buprenorphine gel pack for pain relief. Dams were monitored daily until giving birth to live pups, which were collected for analysis at P10.

### Histology

Adult transgenic L1-EGFP mice (12-16 weeks) were anesthetized using isoflurane, and perfused intracardially with PBS and 4% PFA. CD1 pups, having been electroporated *in utero* with mouse L1-EGFP plasmids, were euthanized at postnatal day 10 by cervical dislocation. 12-week old CBA×C57BL/6 mice, intended for RNA FISH, were injected intraperitoneally with sodium pentobarbital (50mg/kg), followed by cervical dislocation to ensure euthanasia. All brains were dissected and fixed in PFA for 24h. For cryopreservation, fixed brains were immersed first in 15% sucrose and then 30% sucrose to submersion, and embedded in optimal cutting temperature (OCT) compound and stored at -80°C. Transgenic L1-EGFP brains were sectioned on a cryostat (Leica, settings OT=-20°C, CT=-20°C) at 40µm thickness. Free-floating sections were collected in PBS and stored at 4°C. CBA×C57BL/6 brains were sectioned on a cryostat (Leica, settings OT=-22°C, CT=-22°C) at 30µm thickness. Free-floating sections were collected in cryoprotectant (25% glycerol, 35% ethylene glycol, in PBS) and immediately stored at -20°C.

Tissue processing and immunofluorescent staining with primary and secondary antibodies were carried out as described previously^82^. Primary antibodies and dilutions were as follows: rabbit anti-GFP, 1:500 (Thermo Fisher A11122); chicken anti-GFP, 1:500 (Millipore AB16901); mouse anti-T7, 1:500 (Millipore 69522); rabbit anti-T7, 1:500 (Millipore AB3790); goat anti-tdTomato, 1:1000 (Sicgen T2200); mouse anti-NeuN, 1:250 (Millipore MAB377); guinea pig anti-NeuN, 1:250 (Millipore ABN90), rabbit anti-Gad65/67 (GAD1), 1:500 (Sigma G5163); mouse anti-parvalbumin (PV), 1:2000 (Sigma P3088); rabbit anti-β tubulin III (Tub), 1:500 (Sigma T2200); rabbit anti-MeCP2, 1:500 (Abcam ab2828). Secondary antibodies and dilutions were as follows: donkey anti-guinea pig Dylight 405, 1:200 (Jackson Immunoresearch 706475148); donkey anti-mouse Dylight 405, 1:200 (Jackson Immunoresearch 715475150); donkey anti-chicken Alexa Fluor 488, 1:500 (Jackson Immunoresearch 703546155); donkey anti-rat Alexa Fluor 488, 1:500 (Jackson Immunoresearch 712546150); donkey anti-rabbit Alexa Fluor 488, 1:500 (Thermo Fisher A21206); donkey anti-goat Alexa Fluor 594, 1:500 (Jackson Immunoresearch 705586147); donkey anti-rabbit Cy3, 1:200 (Jackson Immunoresearch 711165152); donkey anti-mouse Cy3, 1:500 (Jackson Immunoresearch 715165150); donkey anti-guinea pig Alexa Fluor 647, 1:500 (Millipore AP193SA6); donkey anti-mouse Alexa Fluor, 1:500 (Jackson Immunoresearch 715606150). For nuclei labelling: BisBenzimide H33258 (Sigma B2883). Blocking serum: normal donkey serum (Jackson Immunoresearch 017000121).

### Imaging

EGFP^+^ cells were imaged on a Zeiss LSM510 confocal microscope. Acquisition of high magnification, Z-stack images was performed with Zen 2009 software. Images of EGFP, NeuN and PV immunostaining for quantification were taken from hippocampal and adjacent cortical areas using a Zeiss AxioObserver Z1 microscope and Zen 2009 software, equipped with an ApoTome system and a 10× objective. Visualization and imaging of EGFP, NeuN and PV in *in utero* electroporated mice was performed using a Zeiss Plan-Apochromat 20x/0.8 NA air objective and a Plan-Apochromat 40×/1.4 NA oil-immersion objective on a confocal/two-photon laser-scanning microscope (LSM 710, Carl Zeiss Australia) built around an Axio Observer Z1 body and equipped with two internal gallium arsenide phosphide (GaAsP) photomultiplier tubes (PMTs) and three normal PMTs for epi- (descanned) detection and two external GaAsP PMTs for non-descanned detection in two-photon imaging, and controlled by Zeiss Zen Black software. RNA FISH for sections of hippocampus and adjacent cortical areas, as well as MeCP2, NeuN and PV immunostainings were imaged on a spinning-disk confocal system (Marianas; 3I, Inc.) consisting of a Axio Observer Z1 (Carl Zeiss) equipped with a CSU-W1 spinning-disk head (Yokogawa Corporation of America), ORCA-Flash4.0 v2 sCMOS camera (Hamamatsu Photonics), using a 63×/1.4 NA C-Apo objective and a 20×/0.8 NA Plan-Apochromat objective, respectively. All Z-stack spinning-disk confocal image acquisition was performed using SlideBook 6.0 (3I, Inc). PV stereology was performed on an upright Axio Imager Z2 fluorescent microscope (Carl Zeiss) equipped with a motorized stage and Stereo Investigator software (MBF Bioscience). Contours were drawn based on DAPI staining using a 5×/0.16 NA objective. Counting was performed on a 10×/0.3 NA objective. All image processing and analysis post acquisition were performed using Fiji for Windows (ImageJ 1.52d).

### Single molecule RNA fluorescence *in situ* hybridization (FISH)

Two custom RNAscope probes were designed against the RepBase^83^ L1 T_FI_ subfamily consensus sequence (**Extended Data Fig. 4**). L1 probe A (design #NPR-0003768, Advanced Cell Diagnostics, Cat# ADV827911C3) targeted the L1 T_FI_ 5ʹUTR monomeric and non-monomeric region (consensus positions 827 to 1688). L1 probe B (design #NPR-000412, Advanced Cell Diagnostics, Cat# ADV831481C3) targeted the L1 T_FI_ 5ʹUTR monomeric region (consensus positions 142 to 1423). Weak possible off-target loci for probe A and B comprised the pseudogene Gm-17177, two non-coding RNAs (LOC115486508 for probe A and LOC115490394 for probe B) and a minor isoform of the PPCDC gene (only for probe A), none of which were expressed beyond very low levels or with specificity to PV^+^ neurons. Using the L1 T_F_ RNAscope probes, we performed fluorescence *in situ* hybridization (FISH) on fixed, frozen brain tissue according to the manufacturer’s specifications (RNAscope Fluorescent Multiplex Reagent Kit part 2, Advanced Cell Diagnostics, Cat# 320850) and with the following modifications: 30μm coronal sections instead of 15μm, and boiling in target retrieval solution for 10min instead of 5min. To identify neurons, we performed immunohistochemistry using a rabbit anti-β-tubulin antibody (Sigma Cat# T2200) and donkey anti-rabbit Cy3 secondary antibody (Jackson Immunoresearch, Cat# 711165152) following a previously described protocol^82^. To identify PV^+^ neurons we employed a validated mouse PV RNAscope probe (Mm-Pvalb-C2, Advanced Cell Diagnostics, Cat# ADV421931C2). Probes for the ubiquitously expressed mouse peptidylprolyl isomerase B (PPIB) gene and *Escherichia coli* gene *dapB* were used as positive and negative controls, respectively, for each FISH experiment.

### Cell quantifications

*L1 T_F_* and *PV RNA FISH:* We analyzed four hippocampal sections per animal for each L1 T_F_ 5ʹUTR probe (**Extended Data Fig. 4**) using Imaris 9.5.1 (Bitplane, Oxford Instruments). To render 3D visualizations for a given neuron, we used Tub and DAPI staining to outline its soma and nucleus along Z-stack planes where the cell was detected. We set voxels outside the cell, and inside the nucleus, to a channel intensity value of zero to only retain cytoplasmic L1 mRNA signal and avoid nuclear L1 DNA. We then used the Imaris Spots module to detect L1 mRNA puncta and calculated the number of L1 spots within the cytoplasm in PV^+^/Tub^+^ versus PV^-^/Tub^+^ neurons. *MeCP2*: We quantified MeCP2 protein expression using Imaris 9.5.1 (Bitplane, Oxford Instruments and we analyzed two hippocampal sections per animal. For each cell, we drew the contours of NeuN immunostaining along the relevant Z-stack planes and rendered a cell 3D visualization. We then calculated the mean MeCP2 channel intensity in PV^+^/NeuN^+^ and PV^-^/NeuN^+^ neurons. *PV stereology*: We stained and analyzed every 12th hippocampal section per animal. Cell density was calculated using the total number of PV^+^ cells and the total subregion area from ∼6 sections per animal. *L1-EGFP*: To quantify EGFP^+^ cells we stained and analyzed every 12th hippocampal section (again, ∼6 sections per animal). To visualize colocalization, we used Adobe Photoshop CC 2017. We counted EGFP^+^, EGFP^+^/NeuN^+^ and EGFP^+^/PV^+^ cells across the entire hippocampus and adjacent cortex. The average number of double-labeled cells per 100mm^2^ was determined for each animal. All statistical analyses were performed using Prism (v8.3.1). *ORF1p*: To quantify L1 ORF1p expression, we analyzed two hippocampal sections per animal. We used Imaris 9.5.1 (Bitplane, Oxford Instruments) Surface function to create DG and CA regions of interest and mask selection function to apply the surface to NeuN, ORF1p and PV channels. We then used the spots function (automated cell detection for minimum diameter of 13µm) to identify and quantify NeuN^+^ neurons in each area. We calculated the mean ORF1p and PV channel intensities within each NeuN^+^ neuron. We defined ORF1p^+^ and PV^+^ neurons as those with a mean intensity above a background spot mean intensity.

### Cell sorting and nucleic acid isolation

Neonate litters were obtained from time-mated C57BL/6 mice bred in-house at the QBI animal facility. The day of birth was defined as postnatal day 0 (P0). From each P0 litter of ∼6 pups we dissected and pooled hippocampus tissue. Tissues were dissociated in a papain solution, containing approximately 20U papain (Worthington) and 0.025mg DNase I (Worthington). Prior to use, papain was dissolved in HBSS (Gibco) with 1.1mM EDTA (Invitrogen), 0.067mM mercaptoethanol (Sigma) and 5mM cysteine-HCL (Sigma), and diluted in Hibernate E medium (Gibco). Tissue was incubated for 10min at 37°C with 0.5mL papain solution per embryo. Following digestion, the cell suspension was passed through a 70μm mesh cell strainer, washed into Hibernate E supplemented with B27 (Gibco) and then centrifuged at 300g for 5min. From this point in the protocol onwards, reagents were pre-chilled and the remaining procedures performed on ice. The cell pellet was resuspended in a blocking buffer (HBSS with 5% BSA). A rabbit anti-PV conjugated Alexa Fluor 647 antibody (Bioss bs-1299R-A647, dilution 1:2000) was directly added to the blocking buffer cell suspension and incubated for 1h at 4°C, then passed through a 40μm mesh cell strainer and subjected to flow cytometry. The cell suspension was run through a 100μm nozzle at low pressure (28psi) on a BD FACSAria II flow cytometer (Becton Dickinson). This first sort isolated PV^+^ and PV^-^ cells. To further isolate PV^-^ neurons, PV^-^ cells from the first sort were collected in tubes containing 40U RNAseOUT ribonuclease inhibitor (Invitrogen), then fixed in ice cold 50% ethanol for 5min and centrifuged at 300g for 7min. Following centrifugation, cells were immunostained in blocking buffer containing mouse anti-beta III Tubulin (Tub) conjugated Alexa Fluor 488 antibody (Abcam ab195879, dilution 1:1000) and DAPI (Sigma D9542, 1μg/mL) for 15min at 4°C. Tub^+^ immunostained cells were subjected to a second sort on the same FACS machine and specification as above. Four populations of cells were collected: PV^+^ and PV^-^ (**Extended Data Fig. 6a**, sort 1) and PV^-^/Tub^+^ and PV^-^/Tub^-^ (**Extended Data Fig. 6a**, sort 2). DNA and RNA were then extracted from each cell population. For RNA extractions, cells were sorted directly into the lysis buffer provided in the NucleoSpin RNA XS kit (Macherey Nagel), with RNA extraction performed following the manufacturer’s specifications, except DNAse treatment was performed on a column twice for 20min, instead of once for 15min. For DNA extraction, purified cells were collected into a DNA lysis buffer containing TE buffer (10mM Tris-HCl pH 8 and 0.1mM EDTA), 2% SDS and 100μg/mL proteinase K, and DNA was extracted following a standard phenol-chloroform protocol.

### Quantitative PCR on sorted cells and bulk hippocampus

Total RNA extracted from purified PV^+^, PV^-^, PV^-^/Tub^+^ and PV^-^/Tub^-^ (**Extended Data Fig. 6a**, sorts 1 and 2) populations was used as input for SYBR Green and TaqMan qPCR assays. qPCR reactions were carried out using 300pg RNA/µL from purified PV^+^ and PV^-^ cells and 100pg RNA/µL from purified PV^-^/Tub^+^ and PV^-^/Tub^-^ cells. An RNA integrity number (RIN) above 6, as measured on an Agilent Bioanalyzer (Agilent Technologies, RNA 6000 Pico Kit, Cat# 5067-1513), was set as the minimum cutoff for RNA quality. All qPCRs were carried out on a LightCycler 480 Real-Time PCR system (Roche Life Science). Oligonucleotide PCR primers, as listed in **Supplementary Table 4**, were purchased from Integrated DNA Technologies. *SYBR Green assay*: PCR reactions were prepared using the Power SYBR Green RNA-to-CT 1 step kit (Applied Biosystems, Cat# 4391112). Reactions contained a 2× Power SYBR Green RT-PCR Mix, 10pmol of each primer, 1µL RNA input template and 1× reverse transcriptase enzyme mix in a 10μL final volume. Cycling conditions were as follows: 48°C for 30min, 95°C for 10min, followed by 40 cycles of 95°C, 15sec; 60°C, 1min. To assess potential DNA contamination, an L1 T_F_ qPCR using primers L1Md_5UTR_F and L1Md_5UTR_R was performed with and without reverse transcriptase. A three or more cycle difference between experiments run with and without reverse transcriptase, and detection after cycle 30 in the latter, was considered as non-DNA contaminated RNA. *TaqMan assay*: Applied Biosystems custom L1, URR1 and 5S rRNA TaqMan MGB probes, as listed in **Supplementary Table 4**, were purchased from Thermo Fisher (Cat# 4316032), as was a proprietary mouse GAPDH combination (Cat# 4352339E). TaqMan qPCR reactions contained: 4× TaqPath 1-Step RT-qPCR multiplex reaction master mix (ThermoFisher, Cat# A28521), 4pmol of each primer, 1pmol probe (with the exception of the ORF2/URR1 TaqMan reaction, for which we used 1pmol ORF2 primers) and 1µL RNA input template in a 10uL final volume. Cycling conditions were as follows: 37°C for 2min; 50°C for 15min; 95°C for 2min, followed by 40 cycles of 95°C, 3sec; 60°C, 30sec. TaqMan assays for L1 were multiplexed with assays for either 5S rRNA, GAPDH or URR1 controls. L1 probes were conjugated to VIC or 6FAM fluorophores. Controls were conjugated to HEX, VIC or 6FAM fluorophores. Primer/probe sequences and the associated detection channels are listed in **Supplementary Table 4**. For each assay, the relative mRNA expression in a particular sample was calculated by the delta delta-CT method, using the negative population in the respective sort as control, i.e. PV^+^ was compared to PV^-^ (**Extended Data Fig. 6a**, sort 1) and PV^-^/Tub^+^ compared to PV^-^/Tub^-^ (**Extended Data Fig. 6a**, sort 2). As the PV^-^/Tub^+^ and PV^-^/Tub^-^ populations were isolated as a result of two sortings in serial, for some assays sufficient RNA was only available to perform qPCR on PV^+^ and PV^-^ populations. For qPCR on bulk hippocampus, tissue was isolated from 12-week old animals housed in standard (STD, *N*=12) and enriched (ENR, *N*=14) environments. RNA extraction was performed by Trizol following the manufacturer’s specifications (Trizol reagent, Invitrogen Cat# 15596026). Quantitative TaqMan PCR assays were performed as described above, using 40ng of RNA as input.

### CAPS2.L1 5′RACE and RT-PCR

For 5′RACE, hippocampus tissue from three adult C57BL/6 mice was pooled and RNA extracted (Trizol reagent, Invitrogen Cat# 15596026). RNA was used as input for a FirstChoice RLM-RACE Kit (Invitrogen, Cat# AM1700) to generate cDNA from capped, full-length mRNAs, following the manufacturer’s specifications. Total RNA extracted from purified PV^+^, PV^-^ and pooled neonate hippocampi was reverse transcribed using a High-Capacity cDNA Reverse Transcription Kit (Invitrogen, Cat# 4368814). PCR amplification was then performed using 1U MyTaq HS DNA Polymerase (BioLine) in 1× MyTaq buffer, 10pmol primer CAPS2.L1_F (**Supplementary Table 4**), 10pmol primer CAPS2.L1_R, 1µL cDNA in a 20μL final volume reaction. PCR cycling conditions were as follows: 95°C for 1min, (95°C for 15sec; 55°C for 15sec; 72°C for 10sec)×38, 72°C for 5min. Reaction products were run on a 1.5% agarose gel in 1×TAE, stained with SYBR Safe DNA gel stain.

### L1 T_F_ promoter bisulfite sequencing

Targeted bisulfite sequencing was performed as described previously^45^ to assess L1 T_F_ 5ʹUTR monomer CpG methylation genome-wide. Briefly, this involved extraction of genomic DNA from PV^+^, PV^-^ and PV^-^/Tub^+^ populations purified from hippocampus tissue pooled from neonate littermates (**Extended Data Fig. 6**). Approximately 4×10^4^ events per population were obtained from each of 3 litters (experimental triplicates). DNA was extracted via a conventional phenol-chloroform method and ethanol precipitation aided by glycogen (Ambion). DNA concentration was assessed with a Qubit dsDNA HS assay kit. Next, 20ng of genomic DNA was bisulfite converted using the EZ-DNA Methylation Lightning kit (Zymo Research, Cat# D5030) following the manufacturer’s specifications. Bisulfite PCR reactions used MyTaq HS DNA polymerase (Bioline), and contained 1× reaction buffer, 12.5pmol of each primer, 2µL bisulfite treated DNA input template and 1U of enzyme in a 25µL final volume. PCR cycling conditions were as follows: 95°C for 2min, followed by 40 cycles of 95°C, 30sec; 54°C, 30sec; 72°C, 30sec and 1 cycle of 72°C, 5min. Primer sequences (BS_L1_TF_F and BS_L1_TF_R) were as provided in **Supplementary Table 4**. PCR products were visualized by electrophoresis on a 2% agarose gel, followed by the excision of fragments of expected size and DNA extracted using a MinElute gel extraction kit (Qiagen, Cat# 28604) following the manufacturer’s specifications. DNA concentration was assessed with a Qubit dsDNA HS assay kit and 30ng converted DNA was used as input for library preparation. Libraries were prepared using a NEBNext Ultra II DNA library prep kit (NEB, Cat# E7645S) and NEBNext Multiplex Oligos for Illumina (NEB, Cat# E6609S). Libraries were eluted in 15µL H_2_0 and concentrations measured with an Agilent 2100 Bioanalyzer using an Agilent HS DNA kit (Agilent Technologies, Cat# 5067-4627). Barcoded libraries of PV^+^, PV^-^ and PV^-^/Tub^+^ populations from each of the 3 litters were mixed in equimolar quantities, diluted to 8nM, and combined with 50% PhiX spike-in control (Illumina, Cat# FC-110-3001). Single-end 300mer sequencing was then performed on a MiSeq platform (Illumina) using a MiSeq Reagent v3 kit (Illumina, Cat# MS-102-3003). Data were then analyzed as described elsewhere^6^. To summarize, reads with the L1 T_F_ bisulfite PCR primers at their termini were retained and aligned to the mock converted T_F_ monomer target amplicon sequence with blastn. Reads where non-CpG cytosine bisulfite conversion was <95%, or ≥5% of CpG dinucleotides were mutated, or ≥5% of adenine and guanine nucleotides were mutated, were removed. 100 reads per triplicate cell population, excluding identical bisulfite sequences, were randomly selected and analyzed using QUMA^84^ with default parameters, with strict CpG recognition.

### RNA-seq analysis

The mappability of individual TE copies generally varies as a function of sequencing read length, as well as TE subfamily age and copy number^85,86^. We therefore adopted a prior approach to quantify young mouse (L1 T_F_) and human (L1Hs) subfamily-level transcript abundance with RNA-seq^69,85,87^. Analyzed datasets included Sams *et al*.^40^, bulk hippocampus single-end (1×61mer) RNA-seq obtained from wild-type and conditional CTCF knockout animals (SRA: SRP078142, *N*=3 pools of 2 animals per group), and Yuan *et al*.^39^ bulk single-end (1×49mer) RNA-seq of neurons differentiated *in vitro* from human induced pluripotent stem cells, with and without LHX6 overexpression (SRA: SRP147748, *N*=3 per group). For each RNA-seq library, we aligned reads to the reference genome (mouse: mm10, human: hg38) genome assembly with STAR^88^ version 2.6 (parameters --twopassMode Basic --outSAMprimaryFlag AllBestScore --winAnchorMultimapNmax 1000 --outFilterMultimapNmax 1000) and marked duplicate reads with Picard MarkDuplicates (http://broadinstitute.github.io/picard). We expected the high copy number and limited divergence of young L1 subfamilies to cause most of the corresponding RNA-seq reads to “multi-map” to multiple genomic loci^85,86^. As conceived previously, we assigned multi-map reads a weighting at each of their aligned positions based on the abundance of uniquely mapping reads aligned within 100bp in the same library^69,85,87^. Each position was then assigned a weighting proportionate to the fraction of uniquely mapped reads found there, out of the total number of uniquely mapped reads within 100bp of any mapping position for the multi-mapping read. If no uniquely mapped reads were found near any of the aligned positions for a multi-mapped read, all positions were given an equal weighting. We then intersected the unique and weighted multi-map alignments with RepeatMasker coordinates and produced a total read count for L1 T_F_ (RepeatMasker: “L1Md_T”) and L1Hs genome-wide, normalized by dividing by the total mapped read count for that RNA-seq library (tags-per-million).

### L1 T_F_ 5*′*RACE

Total RNA was extracted from pooled C57BL/6 neonate (postnatal day 1-2) hippocampi and sorted PV^+^ neurons using trizol. 5′RACE was performed using the 5′RACE module of the Takara SMARTer 5′/3′ RACE Kit (Cat# 634858) according to the manufacturer’s protocol. 10ng total RNA was used as input for each reaction, and two reactions from independent experiments were performed. 5′RACE cDNA was PCR amplified using an L1-specific primer (**Supplementary Table 4**) and the 5′RACE universal primer provided in the Takara kit. Thirty-five PCR cycles were performed as follows: 94°C for 30 sec, 67°C for 30 sec, 72°C for 3min. PCR products were visualized on a 1% agarose gel. The resultant smear was excised, and PCR fragments were purified using conventional phenol-chloroform DNA extraction. Iso-Seq template preparation using the Iso-Seq Express kit followed by PacBio SMRT Cell sequencing on a PacBio Sequel II platform was performed by the Australian Genome Research Facility. Four samples (two replicates each of bulk and PV^+^ hippocampus cells) were multiplexed on a single SMRT cell. Reads were aligned to the mouse reference genome (mm10) using minimap2 version 2.20^89^ (parameters -t 96 -N 1000 - p 0.95 -ax splice:hq -uf) and sorted with samtools version 1.12^90^. Uniquely mapped reads, i.e. those that aligned to one genomic position only at their best alignment score and as a primary alignment, were retained if an L1-specific primer and a 5ʹRACE universal primer were located at each of their termini. Reads were then assigned to the full-length L1 T_FI_ elements they overlapped, with alignments required to terminate within the L1 body. The start positions of these alignments within the 5ʹUTR or upstream of the L1 were then recorded as putative transcription start sites (TSSs) supported by at least one read. Replicates gave very similar results and were pooled for display purposes.

### Bulk ATAC-seq analysis

Mouse cortex ATAC-seq data were previously generated by Mo *et al*.^38^ for excitatory pyramidal neurons (marked by CAM2KA), PV interneurons and VIP interneurons, via the isolation of nuclei tagged in specific cell types (INTACT) method. Paired-end fastq files were obtained from the Sequence Read Archive (SRA identifiers SRR1647880-SRR1647885). Trim Galore (parameters --max_n 2 --length 50 --trim-n) was used to apply CutAdapt^91^ to read pairs to trim adapters and low quality bases. Processed reads were aligned to the reference genome (mm10) using bwa mem^92^ with parameters (-a) to output all multimapping alignments. Alignments were filtered to keep only those with an alignment score equal to the maximum achieved for that read. The resulting bam files were sorted using samtools^90^. Peaks for each combined pair of duplicate experiments were called using MACS2^93^ with default parameters, intersected with young L1 genomic coordinates, and then used to calculate the fraction of reads in each replicate aligned to at least one L1-associated peak.

### scATAC-seq analyses

Human hippocampus scATAC-seq data reported by Corces *et al*.^37^ were obtained from the SRA (identifiers SRR11442501 and SRR11442502). Read pairs were retained if the corresponding barcode was present in the 10x Genomics scATAC-seq Unique Molecular Identifier (UMI) whitelist (737K version 1) and then processed and aligned to the hg38 reference genome assembly, as per the bulk ATAC-seq analysis above. Cells (UMIs) with fewer than 10,000 uniquely aligned read pairs were discarded. For the human analysis, a cohort of 277 full-length (>5.9kbp) L1Hs elements defined previously^6^ were employed. Cells were grouped into populations based on having at least one read aligned within the genomic coordinates of the proximal promoter of a given gene, with these coordinates as follows: PV, chr22:36816079-36818079; VIP, chr6:152749797-152751797; GFAP, chr17:44915750-44917750; EXC (CAMK2A), chr5:150289093-150291093. For each cell population, read depth was calculated across each full-length L1Hs copy, and these profiles were then summed to represent the L1Hs subfamily.

### Nanopore methylation analysis

Genomic DNA was phenol-chloroform extracted from PV^+^ and PV^-^ (**Extended Data Fig. 6**) populations purified from 10 neonate (P0) littermate hippocampus sample pools. Additional DNA was obtained from the PV^+^, PV^-^, PV^-^/Tub^+^, PV^-^/Tub^-^ populations of an another neonate pool. DNA samples were sheared to ∼10kb average size, prepared as barcoded libraries using a Ligation Sequencing Kit (Oxford Nanopore Technologies, SQK-LSK109), and sequenced on an ONT PromethION platform (Garvan Sequencing Platform, Australia). Bases were called in SLOW5 format^94^ with Buttery-eel^95^ using Guppy 4.0.11 (Oxford Nanopore Technologies) and reads aligned to the mm10 reference genome using minimap2 version 2.20^89^ and samtools version 1.12^90^. Reads were indexed and per-CpG methylation calls generated using nanopolish version 0.13.2^96^. Methylation likelihood data were sorted by position and indexed using tabix version 1.12^97^. Methylation statistics for the genome divided into 10kbp bins, and reference TEs defined by RepeatMasker coordinates (http://www.repeatmasker.org/), were generated using MethylArtist version 1.0.4^98^, using commands db-nanopolish, segmeth (parameters: --excl_ambig and -- primary_only) and segplot with default parameters. Only full-length (>6kbp) L1s were included, with methylation measured on 5′UTR CpGs. Methylation profiles for individual loci were generated using the MethylArtist command locus, with a 28bp sliding window, and composite profiles with the MethylArtist command composite, with parameters used elsewhere^35^. To identify individual differentially methylated TEs (**Supplementary Table 2**), we required elements to have at least 4 reads and 20 methylation calls in each sample being compared. Comparisons were carried out via Fisher’s Exact Test using methylated and non-methylated call counts, with significance defined as a Bonferroni corrected P value of less than 0.01. Gene ontology enrichment analysis was performed on genes containing at least one differentially methylated L1 T_F_ sequence with PANTHER^51^, using Fisher’s exact test and Bonferroni correction.

### Environmental enrichment and exercise experimental design

At six weeks of age, CBA×C57BL/6 mice were randomly assigned to either a standard (STD), enriched environment (ENR) or exercise (EXE) group, as described previously^99^. All mice were exposed to their assigned housing condition for 6 weeks. Briefly, STD housing consisted of an open-top standard mouse cage (34 × 16 × 16cm; 4 mice/box) with basic bedding and nesting materials. ENR and EXE mice were housed in larger cages (40 × 28 × 18cm; 4 mice/box) containing the same basic bedding and nesting materials as the STD plus specific features. ENR cages contained climbing and tunneling objects together with inanimate objects of various textures, sizes, and shapes, which altogether confer the enhancement of sensory, cognitive and motor stimulation^100^. These cages were changed weekly to ensure novelty for ENR mice. In addition, from weeks 10-12, ENR mice were exposed three times a week for one hour to an extra ‘super-enriched’ condition in a larger playground arena (43 × 80 × 51cm) as previously described^101^. Each EXE cage contained two running wheels (12cm in diameter) to ensure mice had access to voluntary wheel running. Running wheels were excluded from the ENR housing to ensure the effects of physical activity were exclusive to the EXE mice. All mice had *ad libitum* access to food and water and were housed in a controlled room at 22°C and 45% humidity on a 12:12 hour light/dark cycle. All procedures were approved by The Florey Institute of Neuroscience and Mental Health Animal Ethics Committee (19-012-FINMH) and were performed in accordance with the relevant guidelines and regulations of the Australian National Health and Medical Research Council Code of Practice for the Use of Animals for Scientific Purposes.

### CAPS2.L1 expression assays in N2a cells

Differentiating mouse N2a (neuro-2a) neuroblastoma cells (ATCC, Cat# CCL-131) were used to assess the impact of CAPS2.L1 expression on neuronal phenotype. To generate a CAPS2.L1 expression plasmid, we GenScript gene synthesized the CAPS2.L1 chimeric transcript, as detected by ENCODE long-read PacBio sequencing of bulk hippocampus tissue (ENCLB505CBY)^102^ and cloned it to be under the control of a CAG promoter (Vector Builder). To create a same-size control plasmid, we destroyed the *KpnI* restriction site in CAPS2.L1, resulting in *Δ3072* truncation of the CAPS2.L1 ORF. As an additional control, we generated a plasmid expressing the annotated canonical CAPS2.1 transcript (RefSeq: NM_178278.4), also driven by a CAG promoter (Vector Builder). Each plasmid included an EFS promoter driving mCherry to identify cells receiving the constructs. An empty plasmid, containing only EFS driving mCherry expression was used as an additional control. Cells were cultured at 37°C in 5% CO_2_ in Dulbecco’s modified Eagle medium (DMEM) high glucose (4.5g/L), supplemented with 10% FBS and 100U/mL Pen-Strep solution (Life Technologies). 1.2×10^5^ cells per well were plated on coverslips coated with 0.1% gelatine in 12-well plates and transfected with 500ng of plasmid DNA per well using Lipofectamine 2000 (Invitrogen, Cat# 11668019) following the manufacturer’s instructions. 24 hours post-transfection, the medium was replaced with a differentiation medium containing DMEM high glucose with 0.5% FBS and 5µM retinoic acid.

After 5 days of differentiation, culture medium was removed, and cells were fixed in 4% PFA (20min, 4°C) and washed with PBS. Immunohistochemistry was then performed as for mouse primary neuronal cultures. For morphology analysis, Z-stack images were acquired using the SoRa super resolution disk and 3.2× magnification. Image processing and analysis post-acquisition for the morphology analysis were performed using Fiji for Windows (ImageJ 1.52d). NTF3 analysis was performed using Imaris 9.5.1 (Bitplane, Oxford Instruments). To quantify NTF3 release, circular outlines (30µm diameter) were manually added to 3D visualizations of neuronal soma (for mCherry positive neurons) along the Z-stack planes where the cell was detected. Voxels outside the outline were set to a channel intensity value of zero to only retain the NTF3 signal of a given cell. We then manually placed and quantified the number of spots seen as NTF3 signal dots outside of cell soma. For NTF3 analysis inside the soma we used the automatic Imaris surface function to render the surface of mCherry positive cells and exported mean intensities and area. Integrated mean intensity was calculated as equal to area times mean pixel value.

---

**Extended Data Fig. 1:**
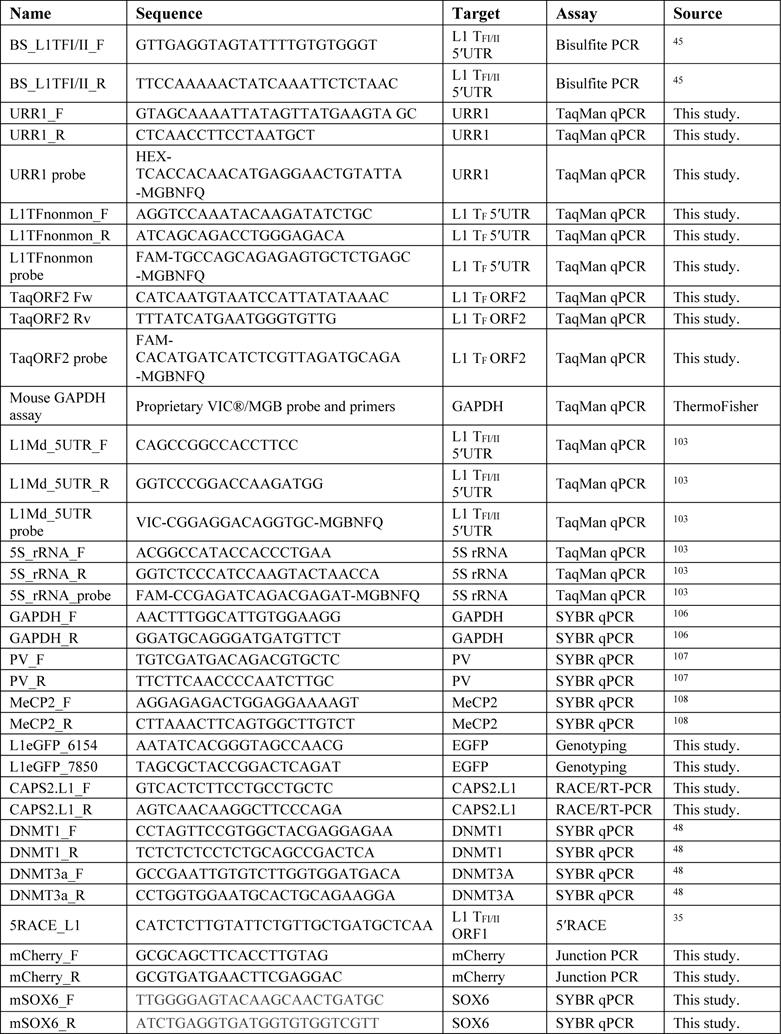
Assessment of L1 retrotransposition in brain and non-brain tissues of L1-EGFP animals. **a**, EGFP cassette genotyping PCR results for the offspring of founder animal #1.2. Circles and squares represent female and male mice, respectively. The indicated 1.7kbp PCR product (red arrow) corresponds to the integrated intron-containing EGFP indicator cassette. Gel labels are as follows: mw, molecular weight (ladder); 1-5, transgenic offspring littermates; +, EGFP positive control plasmid DNA; -, H_2_0. Offspring 2, 4 and 5 carried the L1-EGFP transgene. **b,** Ovary and testis **c,** heart **d,** muscle and **e,** liver tissues of adult L1-EGFP mice were immunostained for EGFP and L1 proteins (T7-tagged ORF1p and 3×FLAG-tagged ORF2p). EGFP^+^ cells were observed in the interstitial cells of the ovaries but not in other tissues. DNA was stained with Hoechst dye (blue). **f,** Representative maximum intensity projection confocal image of a coronal hippocampus section from a transgenic L1-EGFP animal showing immunostaining for EGFP (green) and the interneuron marker GAD1 (red). Yellow arrowheads indicate EGFP^+^/GAD1^+^ neurons in DG. The image is presented as merged and single channels for EGFP and GAD1. Scale bar: 100µm.

**Extended Data Fig. 2:**
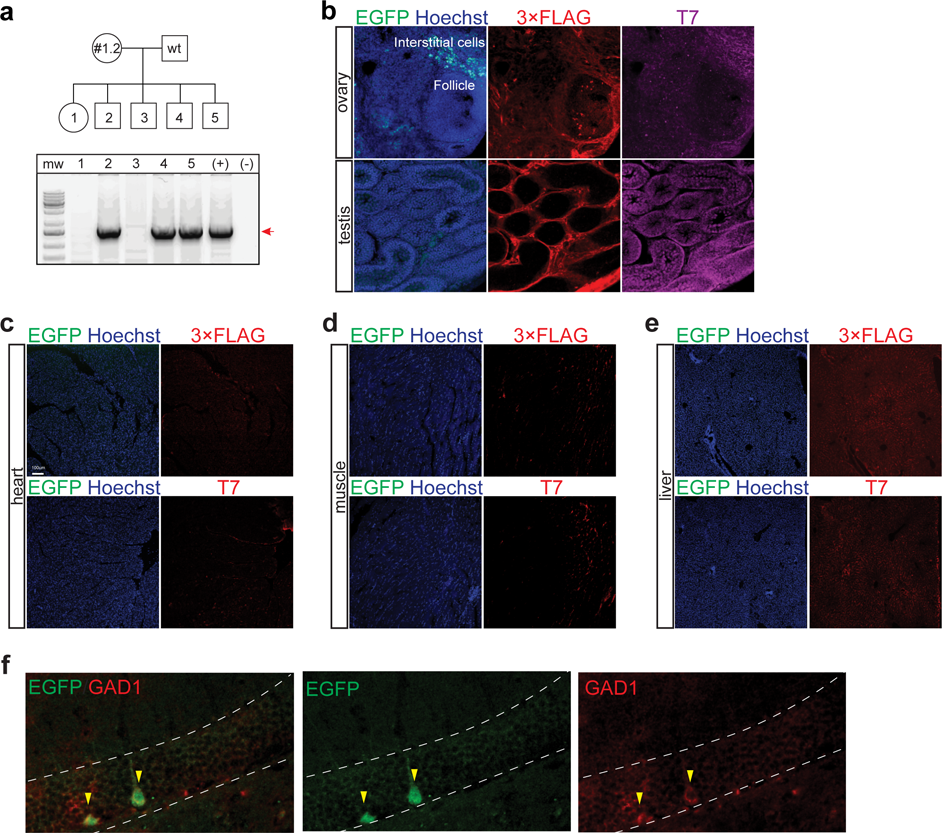
Retrotransposition of an engineered mouse L1 delivered to cultured neurons. **a**, Primary neurons were electroporated with an L1_spa_ reporter construct driven by its native promoter and carrying an mCherry indicator cassette. The mCherry is antisense to the L1, incorporates a β-globin intron in the same orientation as the L1, and is terminated by a polyadenylation signal (filled black lollipop). Retrotransposition removes the β-globin intron, enabling mCherry expression. Experimental timeline is shown below the schematic. Circles represent days in culture, E: electroporation, R: analysis. **b,** Example of a degenerating neuron expressing mCherry (red) and parvalbumin (PV, green). **c,** Representative confocal images of H2A.X (magenta) immunostaining of neurons (Tub, green) shows DNA breaks in mCherry neurons after L1_spa_ electroporation. Immunostaining of L1_spa_ RT^-^ mutant (retrotransposition negative control) and no electroporation controls are shown in the panels below. **d,** Representative confocal images of cL1_spa_ reporter electroporation showing mCherry (red) and SOX6 (magenta) colocalization in primary neurons upon Cre addition. Note: scale bars in (b-d) indicate 10µm.

**Extended Data Fig. 3:**
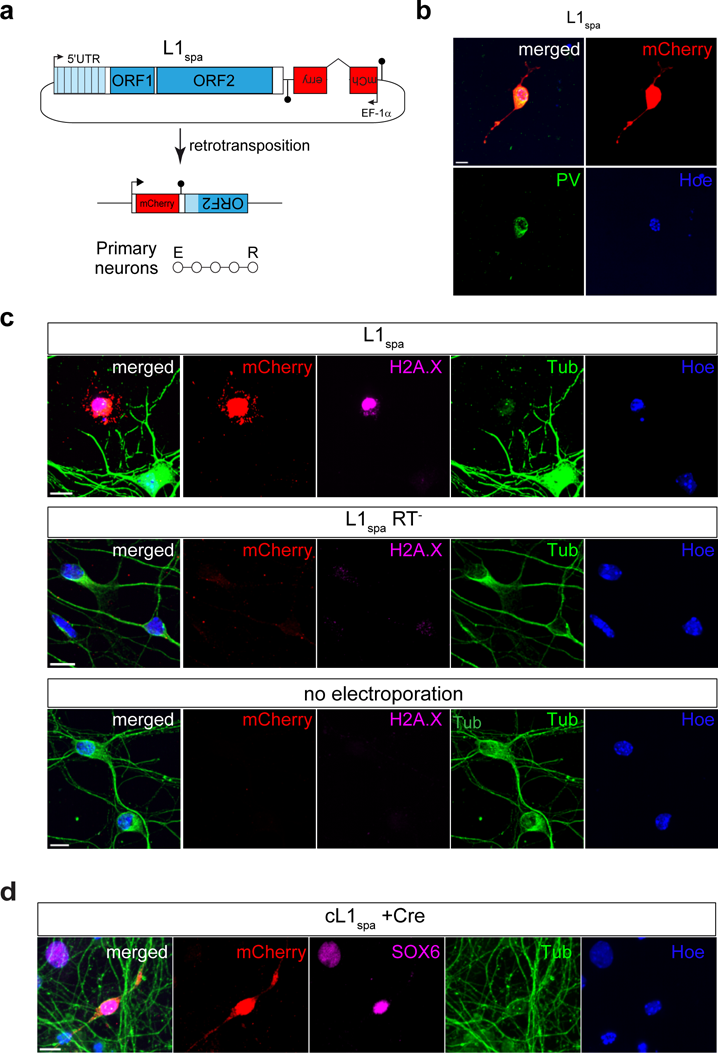
Retrotransposition of an engineered mouse L1 delivered *in utero*. **a**, Schematic representation of L1-EGFP reporter *in utero* electroporation (IUE). A coronal view of electrode positioning is shown at left. Embryos were co-injected with pmCherry (where a CMV promoter controls mCherry expression) and a second plasmid, carrying a mouse L1 tagged with an EGFP indicator cassette (pUBC-L1SM-UBC-EGFP), into the left hemisphere. As a negative control, embryos were co-injected with pmCherry and a disabled L1 reporter plasmid (pMut2-UBC-L1SM-UBC-EGFP) into the right hemisphere. The red inset, shown at right, displays a coronal section of an electroporated mouse brain with pmCherry visible in the targeted hippocampal region. IUE was performed on embryonic day (E)14.5. Embryos were born and then sacrificed at postnatal day (P)10. Note: UBC-L1SM-UBC-EGFP consists of a heterologous UBC promoter driving expression of a synthetic mouse L1 T_F_ (L1SM) containing a native L1 monomeric 5′UTR promoter, codon-optimized ORF1 and ORF2 sequences, the 5′ part of the L1 3′UTR, and an intron-interrupted EGFP indicator cassette with its own UBC promoter and polyadenylation signal (pA). In this system, a cell becomes EGFP positive only when the L1-EGFP mRNA is transcribed, spliced, reverse transcribed and integrated into the genome, allowing a functional EGFP to be expressed from its UBC promoter. Mut2-UBC-L1SM-UBC-EGFP is the same as UBC-L1SM-UBC-EGFP, except it carries mutations known to disable ORF2p reverse transcriptase and endonuclease activities. **b,** Example maximum intensity projection confocal image of a hippocampus section from an embryo electroporated with UBC-L1SM-UBC-EGFP. A PV^+^ (magenta), mCherry^+^ (red), NeuN^+^ (blue), EGFP^+^ cell is indicated with a yellow arrowhead. **c,** No EGFP^+^ cells were found for the retrotransposition-incompetent Mut2-UBC-L1SM-UBC-EGFP plasmid. An empty arrowhead points to a PV^+^, NeuN^+^, mCherry^-^, EGFP^-^ cell.

**Extended Data Fig. 4:**
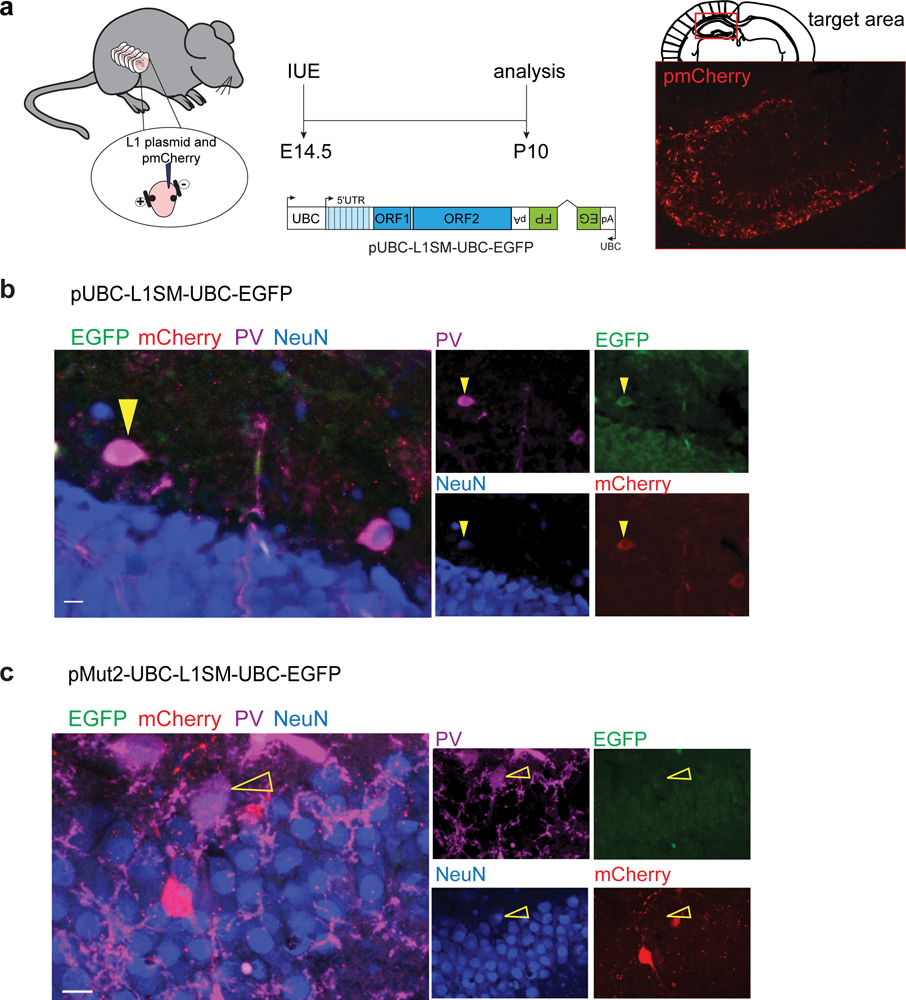
Single-molecule L1 T_F_ RNA FISH experimental design. **a**, L1 T_F_ RNAscope probe A consisted of 20 “ZZ” oligo pairs targeting the L1 T_F_ 5ʹUTR monomeric and non-monomeric region (consensus positions 827 to 1688). **b,** L1 T_F_ probe B was composed of 17 ZZ oligo pairs and targeted the L1 T_F_ 5ʹUTR monomeric region (positions 142 to 1423). **c,** Imaris analysis performed on Z-stack images of L1 T_F_ and PV RNA FISH coronal hippocampus sections immunostained for Tub. Imaris workflow: 1: neuron identification, based on cytoplasmic Tub (red) fluorescence, and cell volume drawing, 2: nucleus definition by DAPI (blue) staining followed by drawing of nuclear volume, 3: delineation of cell and nucleus surfaces, 4: cell surface masking to eliminate voxels outside the cell, 5: nucleus surface masking to exclude voxels inside the nucleus, and 6: L1 T_F_ mRNA (green) fluorescence signal within the defined cytoplasmic volume. **d,** Maximum intensity projection confocal image of a hippocampal section showing L1 T_F_ probe A (green) and PV (magenta) RNA FISH, and Tub (red) immunohistochemistry, in a selected PV^-^ neuron. Dashed lines demark nuclear and cellular boundaries defined for PV and L1 mRNA quantification. **e,** Confocal images of hippocampal sections treated with either DNase or RNase and subsequently processed for RNAscope using the L1 T_F_ probe A (green). **f,** Confocal images of N2a cells transfected with either mouse L1 construct (pL1SM-mCherry) or control (pmCherry) showing specificity of L1 T_F_ RNA FISH probe A (green) signal in cells expressing the L1 construct. **g,** As for (f), except performed on HEK293T cells and showing specificity of L1 T_F_ RNA FISH probe B (green) in cells expressing the L1 construct. **h,** Schematic showing L1 T_F_ sequences assayed by RNA FISH and TaqMan qPCR. Scale bar for (d-g): 10μm.

**Extended Data Fig. 5:**
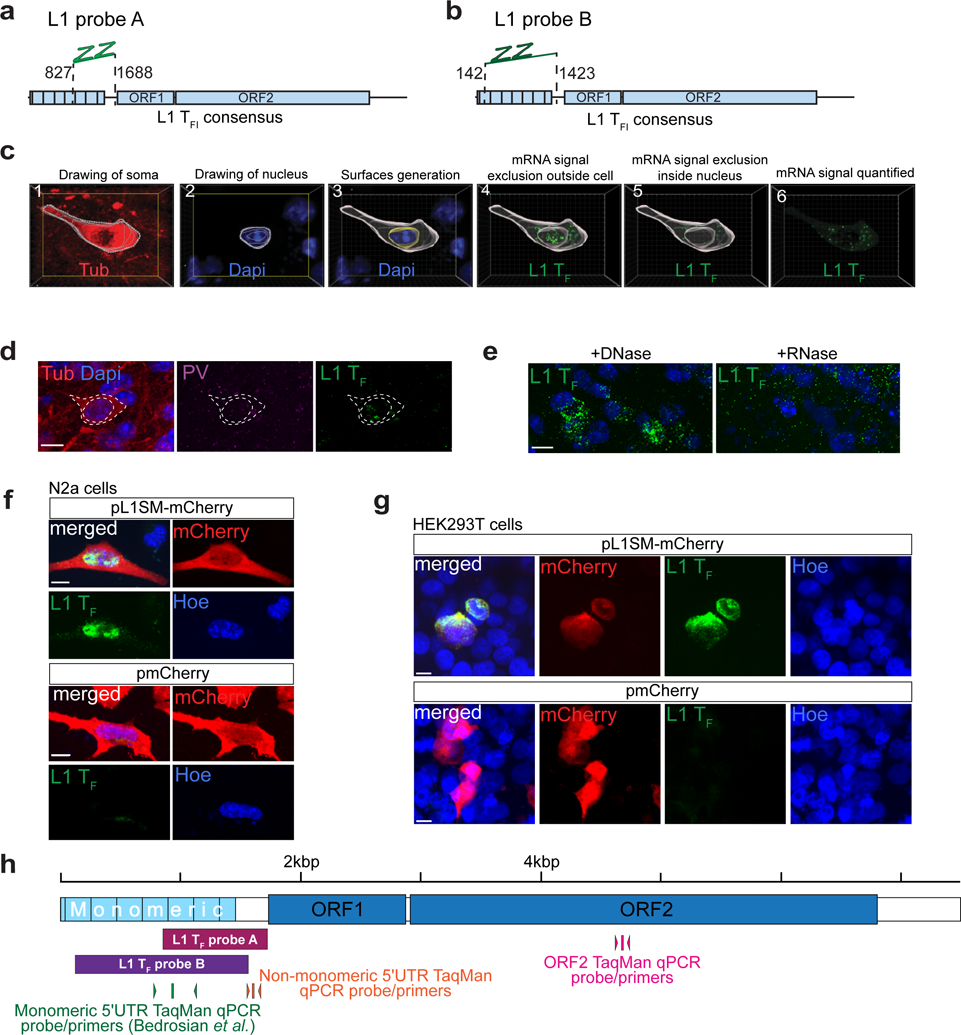
Additional L1 T_F_ RNA FISH data for hippocampus and cortex. **a-c**, Number of L1 T_F_ RNA FISH (probe B) spots in PV^+^/Tub^+^ neurons (orange plot) and PV^-^/Tub^+^ neurons (blue plot) in CA (a), DG (b) and CX (c). \**P*=0.05, \*\**P*=0.01, two-tailed t test comparing animal means, *n*(cells)=5-10, *N*(mice)=3.

**Extended Data Fig. 6:**
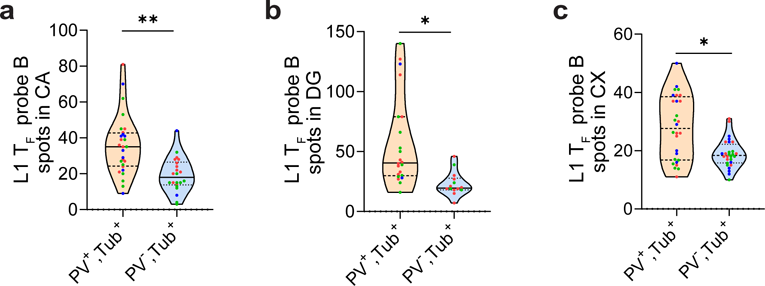
PV^+^ neuron isolation. **a**, Schematic of PV^+^ and PV^-^/Tub^+^ neuron isolation from pooled neonate (P0) litter hippocampal tissue. An anti-PV conjugated antibody (AF647) was used to label and isolate PV^+^ cells. Freshly sorted PV^-^ cells were subsequently labeled with an anti-Tub conjugated antibody (AF488) and sorted again to isolate PV^-^/Tub^+^ neurons. **b,** Gating strategy and purity of fluorescence activated cell sorting (FACS) for PV^+^ cells. **c,** As for (b), except showing PV^-^/Tub^+^ neurons. **d,** Quality control qPCR of relative PV mRNA expression in sorted cells. PV mRNA enrichment in the PV^+^ population was observed, as expected. Data are relative to GAPDH and presented as mean ± SD. \**P*=0.017, two-tailed t test, *N*(litters)=3.

**Extended Data Fig. 7:**
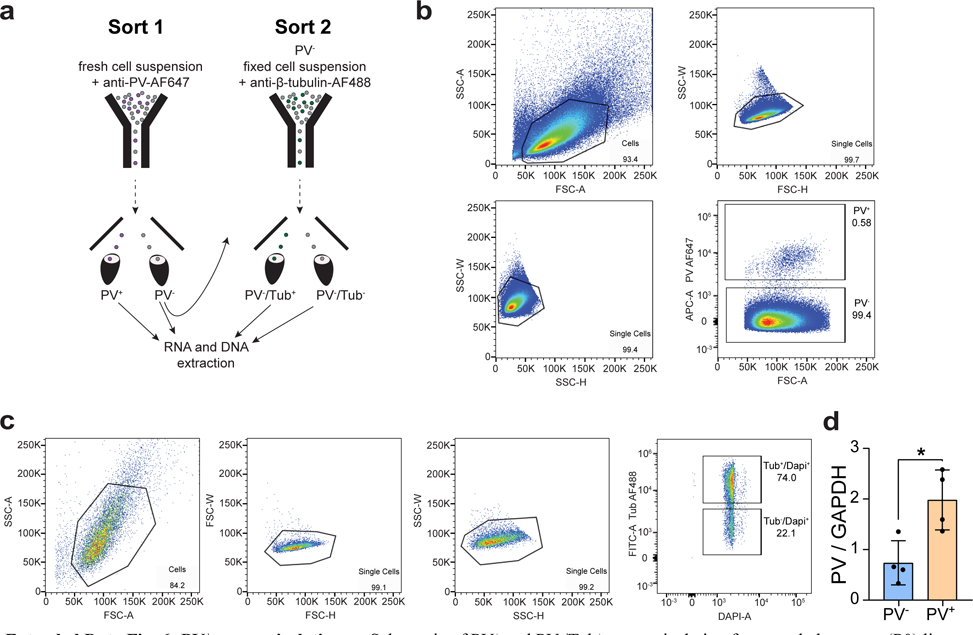
Elevated L1 transcription in PV^+^ neurons. **a**, Multiplexed TaqMan qPCR measuring mRNA abundance of the L1 T_F_ non-monomeric 5ʹUTR sequence (FAM channel) relative to GAPDH (VIC channel) in sorted PV^+^, PV^-^, PV^-^/Tub^+^ and PV^-^/Tub^-^ cells. Cells were sorted from pooled neonate litter hippocampi. *N*=4 litters. **b,** As for (a), except relative to URR1 repetitive DNA (HEX channel). \**P*=0.015. **c,** As for (a), except measuring L1 T_F_ ORF2 (FAM channel) relative to GAPDH (VIC channel). Data are represented as mean ± SD. Significance values were calculated via one-way ANOVA with Tukey’s post-hoc test.

**Extended Data Fig. 8:**
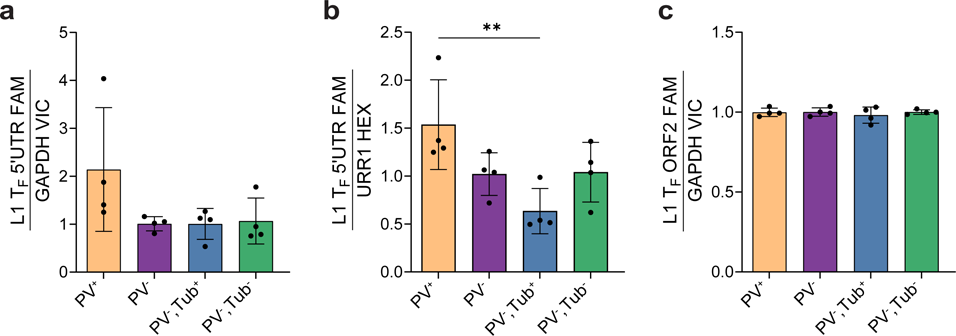
ORF1p antibody specificity. Confocal images of HeLa cells transfected with either a HA-tagged ORF1p (L1_spa_) expression vector or with transfection reagents alone (mock), showing specificity of the ORF1p antibody (green) in human cells expressing mouse ORF1p. Scale bar 10μm.

**Extended Data Fig. 9:**
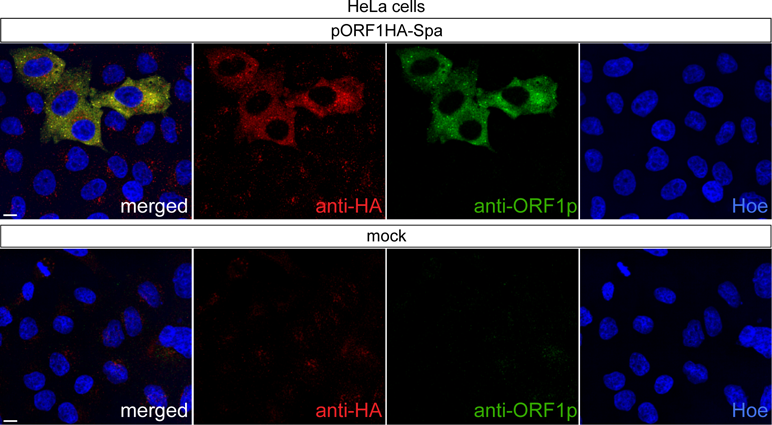
Environmental enrichment does not impact L1 mRNA abundance in PV^+^ neurons. **a**, Standard (STD), exercise (EXE) and enriched (ENR) environment housing schematics. Mice (aged 6 weeks) were placed in either STD, EXE or ENR housing for 6 weeks. ENR and EXE housing consisted of a larger cage with nesting materials. EXE housing contained two running wheels to guarantee all mice had access to voluntary wheel running (excluded in the ENR group). ENR mice were also exposed to spatial stimuli; ladders, tunneling objects and toys of various textures, sizes, and shapes for sensory, cognitive and motor stimulation. Between week 10 and 12, ENR mice were exposed three times a week for one hour to ‘super-enriched’ condition in a larger playground arena with novel toys**. b,** L1 T_F_ RNA FISH spots (probe A) in PV^+^/Tub^+^neurons from STD, EXE and ENR animal CA tissue. *N*(mice)=3-4. Cells from each mouse are color coded. **c,** As for (b) but in DG. **d,** As per (b), except showing L1 T_F_ RNA FISH spots in PV^-^/Tub^+^ neurons. **e,** As for (d), except in DG. **f,** TaqMan qPCR measuring abundance of the L1 T_F_ mRNA monomeric 5ʹUTR (VIC channel) relative to 5S rRNA (FAM channel) in bulk hippocampus samples from STD, EXE and ENR mice. STD *N*=12, ENR *N*=14. **g,** As for (f), except targeting the L1 T_F_ non-monomeric 5ʹUTR (FAM channel) relative to GAPDH (VIC channel). **h,** As for (g), except measuring L1 T_F_ non-monomeric 5ʹUTR (FAM channel) relative to URR1 (HEX channel). **i,** As for (h) except targeting L1 T_F_ ORF2 (FAM channel) relative to URR1 (HEX channel). STD *N*=9, ENR *N*=8. **j,** PV mRNA expression in STD, EXE and ENR conditions, relative to GAPDH. Note: values in (b-j) are represented as mean ± SD. Significance testing was via one-way ANOVA with Tukey’s post-hoc test comparing means of animals. No significant (*P*<0.05) differences were detected between groups.

**Extended Data Fig. 10:**
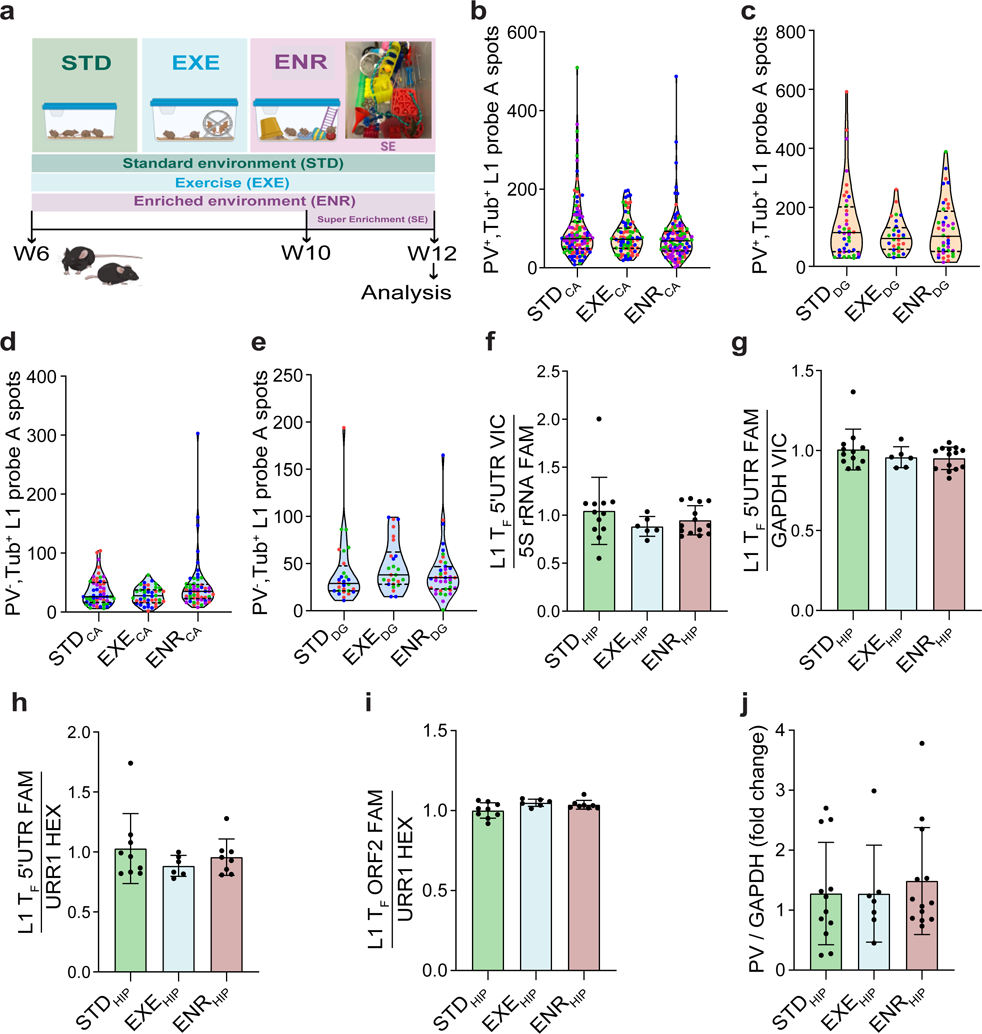
Environmental enrichment does not impact PV^+^ neuron count or L1 ORF1p expression. **a,** Stereological estimation of PV^+^ neuron number in CA of STD, EXE and ENR mice. *N*=3-4 mice per condition. **b,** As per (a), except for DG. **c,** Percentage of L1 ORF1p^+^/PV^+^ versus ORF1p^+^/PV^-^ neurons in STD, EXE and ENR mice. *N*=3-4 mice. **d,** As per (c), except for DG. Note: in (a) and (b) significance testing was via one-way ANOVA and in (c) and (d) via two-way ANOVA, each with Tukey’s post-hoc test. No significant (*P*<0.05) differences were detected between groups.

**Extended Data Fig. 11:**
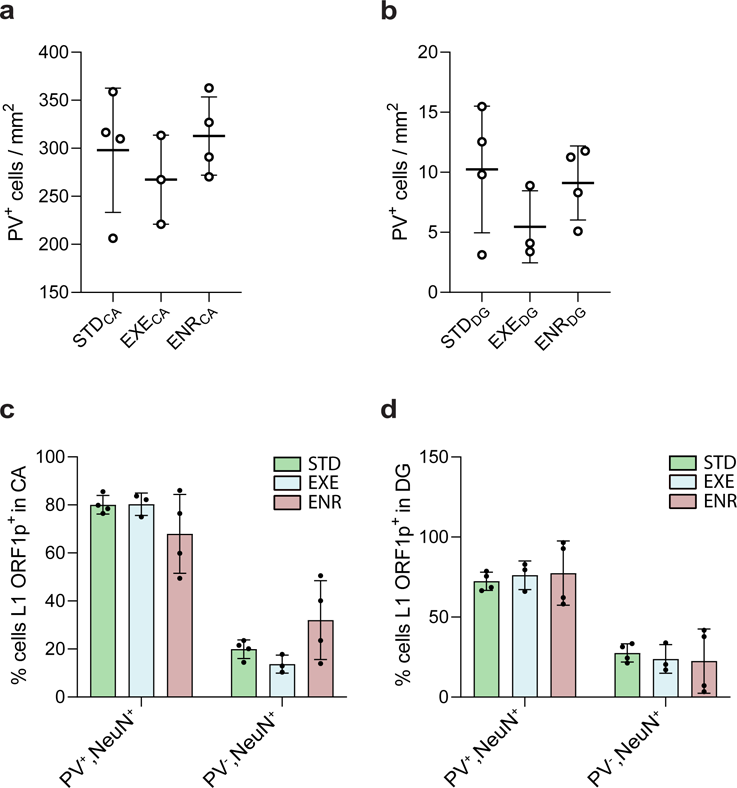
L1 activation by the LHX6/SOX6 transcriptional program. **a**, The L1Hs 5ʹUTR, with YY1- (orange) and SOX-binding (purple) sites marked, is shown above MapRRCon^36^ profiles of ENCODE K562 SOX6 and YY1 ChIP-seq binding and chromatin accessibility profiles in human hippocampal tissue as measured by scATAC-seq^37^. Cells were grouped based on selected accessible genes to define neural cell populations (PV: parvalbumin inhibitory interneuron, EXC: excitatory neurons, VIP: vasoactive intestinal polypeptide-expressing inhibitory interneurons, GFAP: glia). **b,** LHX6, SOX2, SOX5, SOX6, SOX11 and CAPS2 expression in excitatory (EXC) pyramidal neuron, PV interneuron and VIP interneuron cortex populations defined by Mo *et al*.^38^, measured by RNA-seq tags per million (TPM). *N*=2. **c,** Proportion of ATAC-seq reads aligned to peaks associated with full-length L1 T_F_ copies in neuronal populations defined by Mo *et al*.. \*\**P<*0.01, one-way ANOVA with Tukey’s multiple comparison test, *N*=2. **d,** L1Hs subfamily expression measured by RNA-seq TPM in neurons derived via *in vitro* differentiation of induced pluripotent stem cells^39^, with (LHX6↑) and without (control) LHX6 overexpression. \**P*=0.03, two-tailed t test, *N*=3. **e,** L1 T_F_ family expression measured by RNA-seq TPM in bulk hippocampus^40^ of animals with (CTCF cKO) and without (control) conditional knockout of CTCF and associated induction of LHX6 expression. \*\**P*=0.01, two-tailed t test, *N*=3. Note: histogram data in (b-e) are represented as mean ± SD.

**Extended Data Fig. 12:**
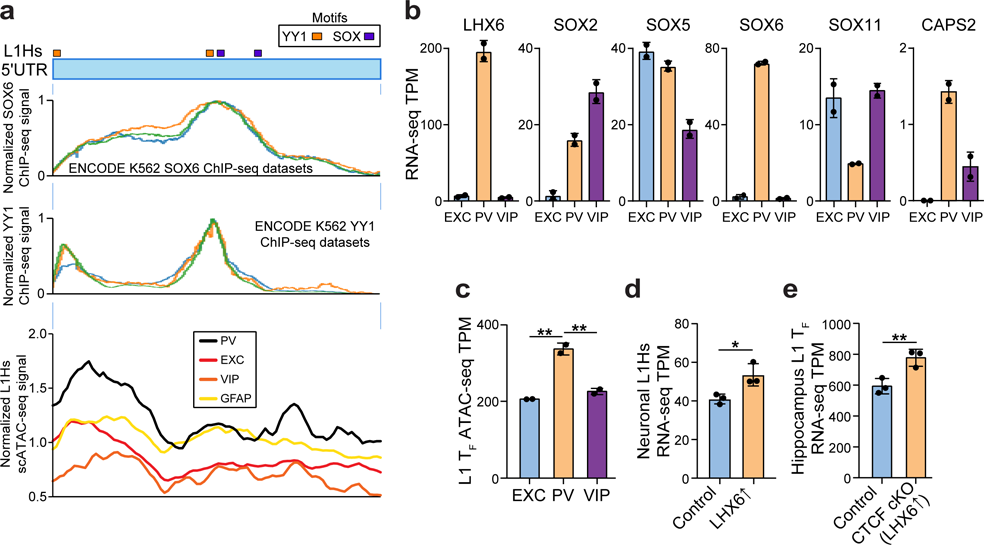
SOX6 overexpression is associated with increased L1 transcription and L1 ORF1p protein expression in primary neuronal cultures. **a**, Confocal images of mouse primary neuronal cultures showing SOX6 (red) expression in Tub^+^ (green) neurons. **b**, Primary neuronal cultures after 5 days *in vitro* were transiently transduced with AAV2 vectors carrying either mCherry or a mouse SOX6 construct (mSOX6-T2A-mCherry) under the control of a CBh promoter. qPCR analysis at 12 and 24h post-transduction revealed a significant increase in SOX6 mRNA expression in mSOX6-T2A-mCherry transduced cultures compared to mCherry controls. \**P*<0.05, two-way ANOVA with Tukey’s multiple comparison test, *N*=4. **c**, TaqMan qPCR measuring abundance of the L1 T_F_ mRNA monomeric 5ʹUTR (VIC channel) relative to 5S rRNA (FAM channel) in mCherry versus mSOX6-T2A-mCherry 24h post-transduction. \**P*=0.012, two-tailed t test, *N*=4. **d**, Images showing L1 ORF1p (magenta), mCherry (red) and Tub (green) immunostaining in neurons 72h post-transduction with AAV2-mCherry. **e**, As per (d), except for mSOX6-T2A-mCherry. **f**, Analysis of L1 ORF1p intensity in mCherry^+^ neurons shows significantly more ORF1p expression in mSOX6-T2A-mCherry transduced neurons than in mCherry controls. \**P*=0.012, two-tailed t test, *N*=3 transductions, *n*=10 cells quantified per experiment. Note: data in (b), (c) and (f) are from independent experiments, represented as mean ± SD. Scale bar: 10μm.

**Extended Data Fig. 13:**
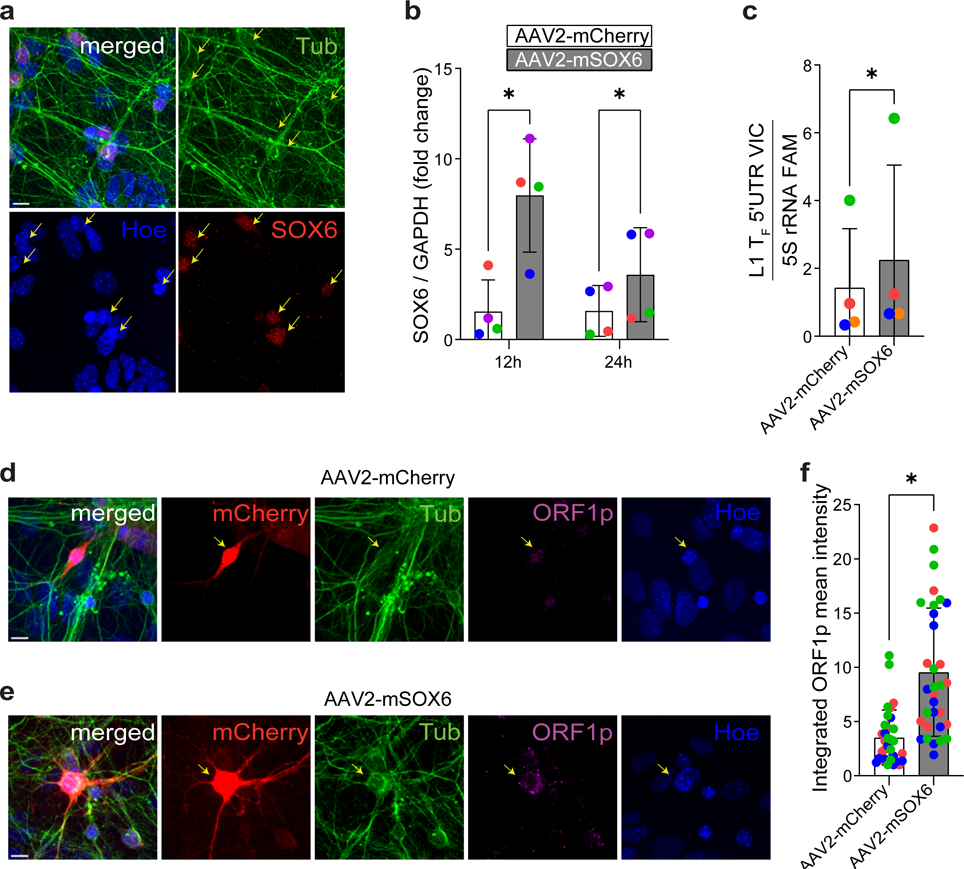
Relaxation of epigenetic repression in PV^+^ neurons. **a**, DNMT1 mRNA abundance measured by qPCR in PV^-^/Tub^+^ and PV^+^ neurons, relative to GAPDH. \**P*=0.038, two-tailed t test, *N*=7 litters. **b,** MeCP2 protein expression in PV^-^/NeuN^+^ (blue plot) and PV^+^/NeuN^+^ (orange plot) neurons. MeCP2 immunofluorescence mean intensities were obtained from coronal hippocampus sections stained for MeCP2, PV and NeuN, and normalized to the PV^-^/NeuN^+^ population mean. \*\*\**P*=0.0007, PV^-^/NeuN^+^ *n*(cells)=414, PV^+^/NeuN^+^ *n*=414, *N*(mice)=4. Cells from each mouse are color coded. **c,** Representative immunostaining image of a coronal hippocampus section showing colocalization of MeCP2 (magenta) with PV (red) and the pan-neuronal marker NeuN (green). Yellow arrows indicate PV^+^ neurons on single channel images. Scale bar: 10μm.

**Extended Data Fig. 14:**
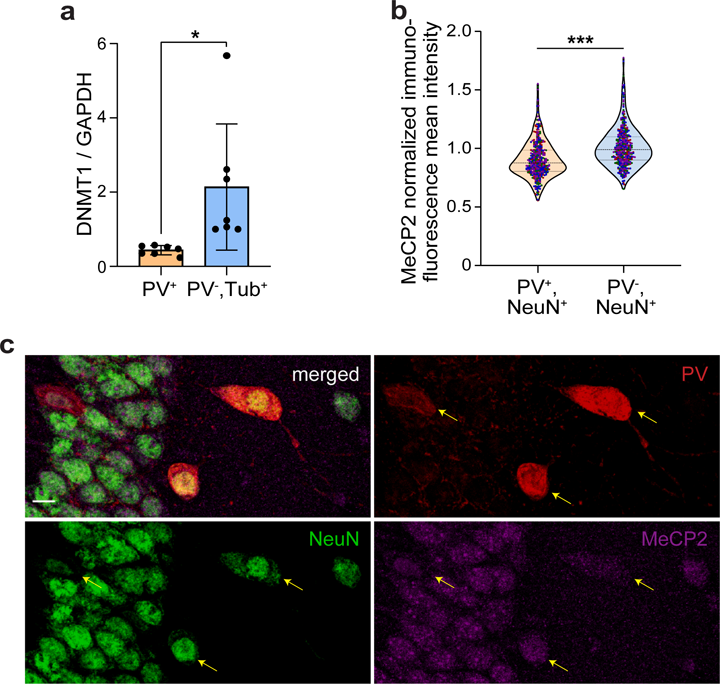
Hypomethylated L1s in PV^+^ neuron genes, as identified by ONT sequencing. **a**, Methylation profile of a full-length L1 T_FIII_ element with intact ORFs, intronic to CHL1. The first panel shows the L1 orientated in sense to the last intron of CHL1. The second panel displays aligned ONT reads, with unmethylated CpGs colored in orange (PV^+^) and purple (PV^-^), blue (PV^-^/Tub^+^) and green (PV^-^/Tub^-^), and methylated CpGs colored black. The third panel indicates the relationship between CpG positions in genome space and CpG space, including those corresponding to the L1 T_FIII_ 5ʹUTR (shaded light green). The fourth panel indicates the fraction of methylated CpGs for each cell type across CpG space. **b,** As for (a), except displaying an L1 T_FIII_ antisense and intronic to ERBB4.

**Extended Data Fig. 15:**
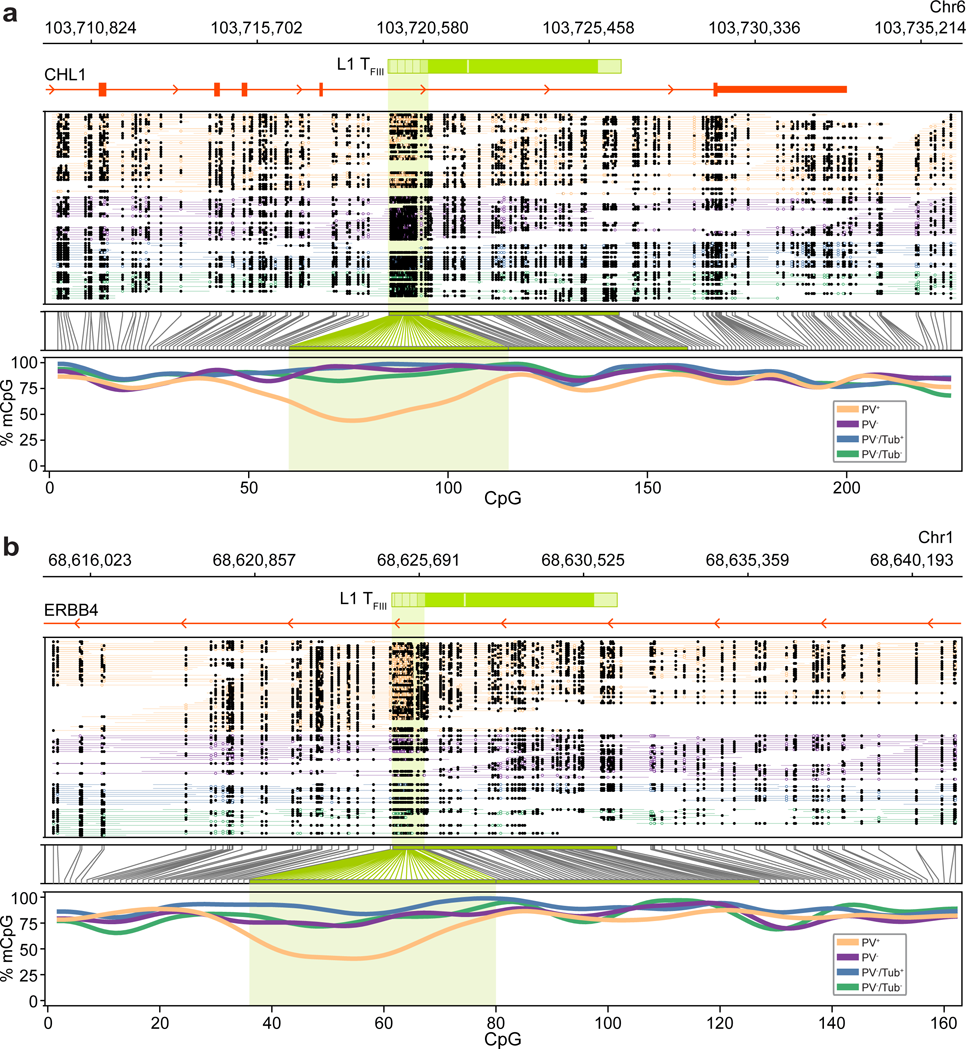
Model of L1 activation by SOX6 in PV^+^ neurons.

**Supplementary Table 4.**
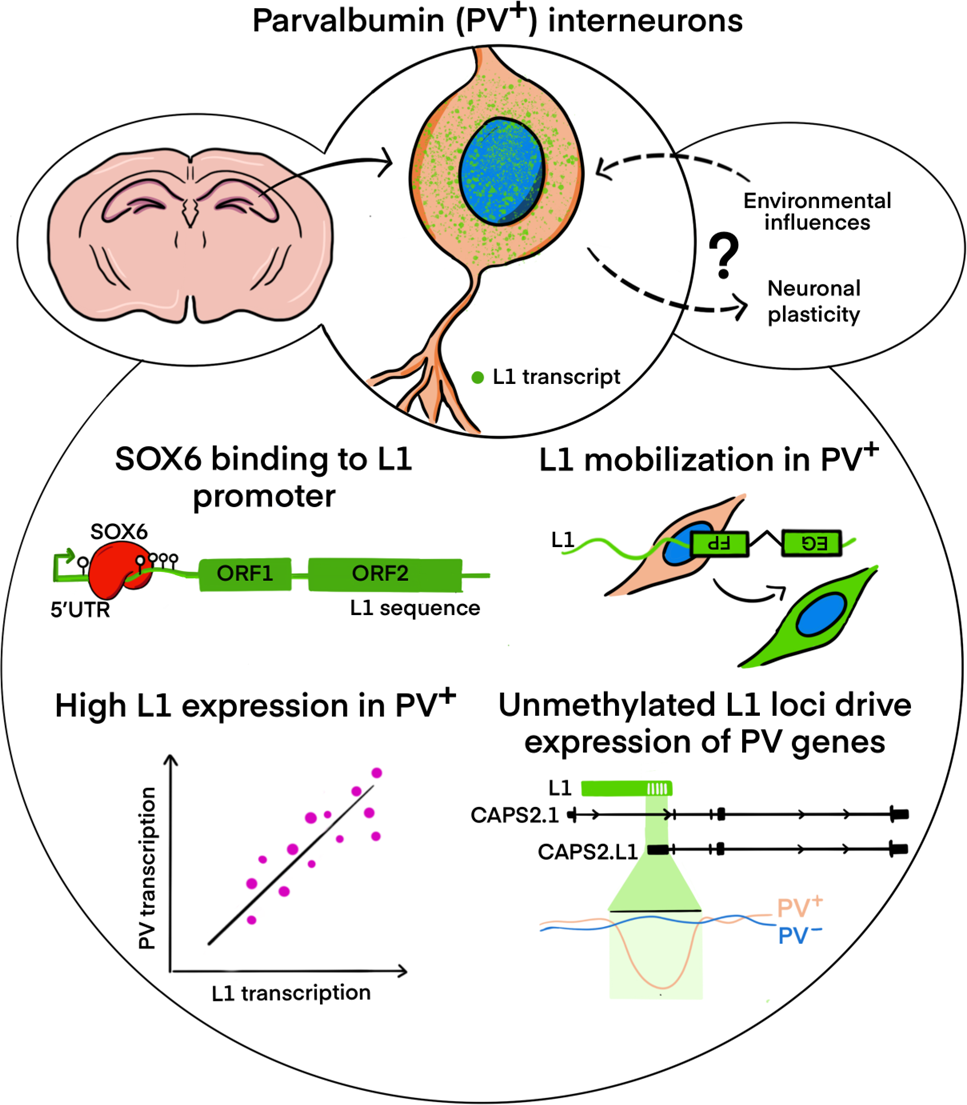
Primer and probe information.

## References

1. Chuong, E. B., Elde, N. C. & Feschotte, C. Regulatory activities of transposable elements: from conflicts to benefits. Nat. Rev. Genet. 18, 71–86 (2017).

2. Britten, R. J. & Davidson, E. H. Gene regulation for higher cells: a theory. Science 165, 349– 357 (1969).

3. Senft, A. D. & Macfarlan, T. S. Transposable elements shape the evolution of mammalian development. Nat. Rev. Genet. 22, 691–711 (2021).

4. Erwin, J. A. et al. L1-associated genomic regions are deleted in somatic cells of the healthy human brain. Nat. Neurosci. 19, 1583–1591 (2016).

5. Evrony, G. D. et al. Cell lineage analysis in human brain using endogenous retroelements. Neuron 85, 49–59 (2015).

6. Sanchez-Luque, F. J. et al. LINE-1 Evasion of Epigenetic Repression in Humans. Mol. Cell 75, 590–604 (2019).

7. Billon, V. et al. Somatic retrotransposition in the developing rhesus macaque brain. Genome Res. 32, 1298–1314 (2022).

8. Batista-Brito, R. et al. The cell-intrinsic requirement of Sox6 for cortical interneuron development. Neuron 63, 466–481 (2009).

9. Connor, F., Wright, E., Denny, P., Koopman, P. & Ashworth, A. The Sry-related HMG box-containing gene Sox6 is expressed in the adult testis and developing nervous system of the mouse. Nucleic Acids Res. 23, 3365–3372 (1995).

10. Munguba, H. et al. Postnatal Sox6 Regulates Synaptic Function of Cortical Parvalbumin-Expressing Neurons. J. Neurosci. 41, 8876–8886 (2021).

11. Kazazian, H. H., Jr & Moran, J. V. Mobile DNA in Health and Disease. N. Engl. J. Med. 377, 361–370 (2017).

12. Moran, J. V., DeBerardinis, R. J. & Kazazian, H. H., Jr. Exon shuffling by L1 retrotransposition. Science 283, 1530–1534 (1999).

13. Denli, A. M. et al. Primate-specific ORF0 contributes to retrotransposon-mediated diversity. Cell 163, 583–593 (2015).

14. Moran, J. V. et al. High frequency retrotransposition in cultured mammalian cells. Cell 87, 917–927 (1996).

15. Wei, W. et al. Human L1 retrotransposition: cis preference versus trans complementation. Mol. Cell. Biol. 21, 1429–1439 (2001).

16. Feng, Q., Moran, J. V., Kazazian, H. H., Jr & Boeke, J. D. Human L1 retrotransposon encodes a conserved endonuclease required for retrotransposition. Cell 87, 905–916 (1996).

17. Kazazian, H. H., Jr et al. Haemophilia A resulting from de novo insertion of L1 sequences represents a novel mechanism for mutation in man. Nature 332, 164–166 (1988).

18. van den Hurk, J. A. J. M., et al. L1 retrotransposition can occur early in human embryonic development. Hum. Mol. Genet. 16, 1587–1592 (2007).

19. Richardson, S. R. et al. Heritable L1 retrotransposition in the mouse primordial germline and early embryo. Genome Res. 27, 1395–1405 (2017).

20. An, W. et al. Active retrotransposition by a synthetic L1 element in mice. Proc. Natl. Acad. Sci. U. S. A. 103, 18662–18667 (2006).

21. Feusier, J. et al. Pedigree-based estimation of human mobile element retrotransposition rates. Genome Res. 29, 1567–1577 (2019).

22. Muotri, A. R. et al. Somatic mosaicism in neuronal precursor cells mediated by L1 retrotransposition. Nature 435, 903–910 (2005).

23. Coufal, N. G. et al. L1 retrotransposition in human neural progenitor cells. Nature 460, 1127– 1131 (2009).

24. Ostertag, E. M., Prak, E. T., DeBerardinis, R. J., Moran, J. V. & Kazazian, H. H., Jr. Determination of L1 retrotransposition kinetics in cultured cells. Nucleic Acids Res. 28, 1418– 1423 (2000).

25. Sassaman, D. M. et al. Many human L1 elements are capable of retrotransposition. Nat. Genet. 16, 37–43 (1997).

26. Dombroski, B. A., Scott, A. F. & Kazazian, H. H., Jr. Two additional potential retrotransposons isolated from a human L1 subfamily that contains an active retrotransposable element. Proc. Natl. Acad. Sci. U. S. A. 90, 6513–6517 (1993).

27. Donato, F., Rompani, S. B. & Caroni, P. Parvalbumin-expressing basket-cell network plasticity induced by experience regulates adult learning. Nature 504, 272–276 (2013).

28. Naas, T. P. et al. An actively retrotransposing, novel subfamily of mouse L1 elements. EMBO J. 17, 590–597 (1998).

29. Richardson, S. R. et al. Revisiting the impact of synthetic ORF sequences on engineered LINE-1 retrotransposition. bioRxiv 2022.08.29.505632 (2022) doi:10.1101/2022.08.29.505632.

30. Gerdes, P. et al. Retrotransposon instability dominates the acquired mutation landscape of mouse induced pluripotent stem cells. Nat. Commun. 13, 7470 (2022).

31. Han, J. S. & Boeke, J. D. A highly active synthetic mammalian retrotransposon. Nature 429, 314–318 (2004).

32. DeBerardinis, R. J. & Kazazian, H. H., Jr. Analysis of the promoter from an expanding mouse retrotransposon subfamily. Genomics 56, 317–323 (1999).

33. Athanikar, J. N., Badge, R. M. & Moran, J. V. A YY1-binding site is required for accurate human LINE-1 transcription initiation. Nucleic Acids Res. 32, 3846–3855 (2004).

34. Tchénio, T., Casella, J. F. & Heidmann, T. Members of the SRY family regulate the human LINE retrotransposons. Nucleic Acids Res. 28, 411–415 (2000).

35. Gerdes, P. et al. Locus-resolution analysis of L1 regulation and retrotransposition potential in mouse embryonic development. Genome Res. 33, 1465–1481 (2023).

36. Sun, X. et al. Transcription factor profiling reveals molecular choreography and key regulators of human retrotransposon expression. Proc. Natl. Acad. Sci. U. S. A. 115, E5526– E5535 (2018).

37. Corces, M. R. et al. Single-cell epigenomic analyses implicate candidate causal variants at inherited risk loci for Alzheimer’s and Parkinson’s diseases. Nat. Genet. 52, 1158–1168 (2020).

38. Mo, A. et al. Epigenomic Signatures of Neuronal Diversity in the Mammalian Brain. Neuron 86, 1369–1384 (2015).

39. Yuan, F. et al. Induction of human somatostatin and parvalbumin neurons by expressing a single transcription factor LIM homeobox 6. Elife 7, e37382 (2018).

40. Sams, D. S. et al. Neuronal CTCF Is Necessary for Basal and Experience-Dependent Gene Regulation, Memory Formation, and Genomic Structure of BDNF and Arc. Cell Rep. 17, 2418–2430 (2016).

41. Brouha, B. et al. Evidence consistent with human L1 retrotransposition in maternal meiosis I. Am. J. Hum. Genet. 71, 327–336 (2002).

42. Kopera, H. C. et al. LINE-1 cultured cell retrotransposition assay. Methods Mol. Biol. 1400, 139–156 (2016).

43. Castro-Diaz, N. et al. Evolutionally dynamic L1 regulation in embryonic stem cells. Genes Dev. 28, 1397–1409 (2014).

44. de la Rica, L. et al. TET-dependent regulation of retrotransposable elements in mouse embryonic stem cells. Genome Biol. 17, 234 (2016).

45. Schauer, S. N. et al. L1 retrotransposition is a common feature of mammalian hepatocarcinogenesis. Genome Res. 28, 639–653 (2018).

46. Lavery, L. A. et al. Losing Dnmt3a dependent methylation in inhibitory neurons impairs neural function by a mechanism impacting Rett syndrome. Elife 9, e52981 (2020).

47. Wu, H. et al. Dnmt3a-dependent nonpromoter DNA methylation facilitates transcription of neurogenic genes. Science 329, 444–448 (2010).

48. Feng, J. et al. Dnmt1 and Dnmt3a maintain DNA methylation and regulate synaptic function in adult forebrain neurons. Nat. Neurosci. 13, 423–430 (2010).

49. Muotri, A. R. et al. L1 retrotransposition in neurons is modulated by MeCP2. Nature 468, 443–446 (2010).

50. Ewing, A. D. et al. Nanopore Sequencing Enables Comprehensive Transposable Element Epigenomic Profiling. Mol. Cell 80, 915–928.e5 (2020).

51. Thomas, P. D. et al. PANTHER: Making genome-scale phylogenetics accessible to all. Protein Sci. 31, 8–22 (2022).

52. Sookdeo, A., Hepp, C. M., McClure, M. A. & Boissinot, S. Revisiting the evolution of mouse LINE-1 in the genomic era. Mob. DNA 4, 3 (2013).

53. Girard, F., Venail, J., Schwaller, B. & Celio, M. R. The EF-hand Ca2+-binding protein super-family: A genome-wide analysis of gene expression patterns in the adult mouse brain. Neuroscience 294, 116–155 (2015).

54. Schmalbach, B. et al. Age-dependent loss of parvalbumin-expressing hippocampal interneurons in mice deficient in CHL1, a mental retardation and schizophrenia susceptibility gene. J. Neurochem. 135, 830–844 (2015).

55. Chen, Y.-J. et al. ErbB4 in parvalbumin-positive interneurons is critical for neuregulin 1 regulation of long-term potentiation. Proc. Natl. Acad. Sci. U. S. A. 107, 21818–21823 (2010).

56. Duangdao, D. M., Clark, S. D., Okamura, N. & Reinscheid, R. K. Behavioral phenotyping of neuropeptide S receptor knockout mice. Behav. Brain Res. 205, 1–9 (2009).

57. ENCODE Project Consortium. An integrated encyclopedia of DNA elements in the human genome. Nature 489, 57–74 (2012).

58. Fuchs, E. C. et al. Recruitment of parvalbumin-positive interneurons determines hippocampal function and associated behavior. Neuron 53, 591–604 (2007).

59. Ognjanovski, N. et al. Parvalbumin-expressing interneurons coordinate hippocampal network dynamics required for memory consolidation. Nat. Commun. 8, 15039 (2017).

60. Erwin, J. A., Marchetto, M. C. & Gage, F. H. Mobile DNA elements in the generation of diversity and complexity in the brain. Nat. Rev. Neurosci. 15, 497–506 (2014).

61. McConnell, M. J. et al. Intersection of diverse neuronal genomes and neuropsychiatric disease: The Brain Somatic Mosaicism Network. Science 356, eaal1641 (2017).

62. Muotri, A. R. & Gage, F. H. Generation of neuronal variability and complexity. Nature 441, 1087–1093 (2006).

63. Baillie, J. K. et al. Somatic retrotransposition alters the genetic landscape of the human brain. Nature 479, 534–537 (2011).

64. Miki, Y. et al. Disruption of the APC gene by a retrotransposal insertion of L1 sequence in a colon cancer. Cancer Res. 52, 643–645 (1992).

65. Scott, E. C. et al. A hot L1 retrotransposon evades somatic repression and initiates human colorectal cancer. Genome Res. 26, 745–755 (2016).

66. Howell, R. & Usdin, K. The ability to form intrastrand tetraplexes is an evolutionarily conserved feature of the 3’end of L1 retrotransposons. Mol. Biol. Evol. 14, 144–155 (1997).

67. Deniz, Ö., Frost, J. M. & Branco, M. R. Regulation of transposable elements by DNA modifications. Nat. Rev. Genet. 20, 417–431 (2019).

68. Jönsson, M. E. et al. Activation of neuronal genes via LINE-1 elements upon global DNA demethylation in human neural progenitors. Nat. Commun. 10, 3182 (2019).

69. Faulkner, G. J. et al. The regulated retrotransposon transcriptome of mammalian cells. Nat. Genet. 41, 563–571 (2009).

70. Rose, N. R. & Klose, R. J. Understanding the relationship between DNA methylation and histone lysine methylation. Biochim. Biophys. Acta 1839, 1362–1372 (2014).

71. Bourc’his, D. & Bestor, T. H. Meiotic catastrophe and retrotransposon reactivation in male germ cells lacking Dnmt3L. Nature 431, 96–99 (2004).

72. Ooi, S. K. T. et al. DNMT3L connects unmethylated lysine 4 of histone H3 to de novo methylation of DNA. Nature 448, 714–717 (2007).

73. Garcia-Perez, J. L. et al. Epigenetic silencing of engineered L1 retrotransposition events in human embryonic carcinoma cells. Nature 466, 769–773 (2010).

74. Yusa, K., Zhou, L., Li, M. A., Bradley, A. & Craig, N. L. A hyperactive piggyBac transposase for mammalian applications. Proc. Natl. Acad. Sci. U. S. A. 108, 1531–1536 (2011).

75. Garcia-Perez, J. L. et al. LINE-1 retrotransposition in human embryonic stem cells. Hum. Mol. Genet. 16, 1569–1577 (2007).

76. Alisch, R. S., Garcia-Perez, J. L., Muotri, A. R., Gage, F. H. & Moran, J. V. Unconventional translation of mammalian LINE-1 retrotransposons. Genes Dev. 20, 210–224 (2006).

77. Sauer, B. & Henderson, N. Site-specific DNA recombination in mammalian cells by the Cre recombinase of bacteriophage P1. Proc. Natl. Acad. Sci. U. S. A. 85, 5166–5170 (1988).

78. Sternberg, N. & Hamilton, D. Bacteriophage P1 site-specific recombination. I. Recombination between loxP sites. J. Mol. Biol. 150, 467–486 (1981).

79. An, W. et al. Conditional activation of a single-copy L1 transgene in mice by Cre. Genesis 46, 373–383 (2008).

80. Campbell, B. C. et al. mGreenLantern: a bright monomeric fluorescent protein with rapid expression and cell filling properties for neuronal imaging. Proc. Natl. Acad. Sci. U. S. A. 117, 30710–30721 (2020).

81. Paolino, A. et al. Differential timing of a conserved transcriptional network underlies divergent cortical projection routes across mammalian brain evolution. Proc. Natl. Acad. Sci. U. S. A. 117, 10554–10564 (2020).

82. Blaess, S. et al. Temporal-spatial changes in Sonic Hedgehog expression and signaling reveal different potentials of ventral mesencephalic progenitors to populate distinct ventral midbrain nuclei. Neural Dev. 6, 29 (2011).

83. Bao, W., Kojima, K. K. & Kohany, O. Repbase Update, a database of repetitive elements in eukaryotic genomes. Mob. DNA 6, 11 (2015).

84. Kumaki, Y., Oda, M. & Okano, M. QUMA: quantification tool for methylation analysis. Nucleic Acids Res. 36, W170–5 (2008).

85. Faulkner, G. J. et al. A rescue strategy for multimapping short sequence tags refines surveys of transcriptional activity by CAGE. Genomics 91, 281–288 (2008).

86. Lanciano, S. & Cristofari, G. Measuring and interpreting transposable element expression. Nat. Rev. Genet. 21, 721–736 (2020).

87. Hashimoto, T. et al. Probabilistic resolution of multi-mapping reads in massively parallel sequencing data using MuMRescueLite. Bioinformatics 25, 2613–2614 (2009).

88. Dobin, A. et al. STAR: ultrafast universal RNA-seq aligner. Bioinformatics 29, 15–21 (2013).

89. Li, H. Minimap2: pairwise alignment for nucleotide sequences. Bioinformatics 34, 3094– 3100 (2018).

90. Li, H. et al. The Sequence Alignment/Map format and SAMtools. Bioinformatics 25, 2078–2079 (2009).

91. Martin, M. Cutadapt removes adapter sequences from high-throughput sequencing reads. EMBnet.journal 17, 10–12 (2011).

92. Li, H. Aligning sequence reads, clone sequences and assembly contigs with BWA-MEM. arXiv [q-bio.GN] arXiv:1303.3997 (2013).

93. Zhang, Y. et al. Model-based analysis of ChIP-Seq (MACS). Genome Biol. 9, R137 (2008).

94. Gamaarachchi, H. et al. Fast nanopore sequencing data analysis with SLOW5. Nat. Biotechnol. 40, 1026–1029 (2022).

95. Samarakoon, H., Ferguson, J. M., Gamaarachchi, H. & Deveson, I. W. Accelerated nanopore basecalling with SLOW5 data format. Bioinformatics 39, (2023).

96. Simpson, J. T. et al. Detecting DNA cytosine methylation using nanopore sequencing. Nat. Methods 14, 407–410 (2017).

97. Li, H. Tabix: fast retrieval of sequence features from generic TAB-delimited files. Bioinformatics 27, 718–719 (2011).

98. Cheetham, S. W., Kindlova, M. & Ewing, A. D. Methylartist: Tools for Visualising Modified Bases from Nanopore Sequence Data. Bioinformatics 38, 3109–3112 (2022).

99. Love, C. J., Gubert, C., Renoir, T. & Hannan, A. J. Environmental enrichment and exercise housing protocols for mice. STAR Protoc 3, 101689 (2022).

100. Gubert, C. & Hannan, A. J. Environmental enrichment as an experience-dependent modulator of social plasticity and cognition. Brain Res. 1717, 1–14 (2019).

101. Mazarakis, N. K. et al. ‘Super-Enrichment’ Reveals Dose-Dependent Therapeutic Effects of Environmental Stimulation in a Transgenic Mouse Model of Huntington’s Disease. J. Huntingtons Dis. 3, 299–309 (2014).

102. Luo, Y. et al. New developments on the Encyclopedia of DNA Elements (ENCODE) data portal. Nucleic Acids Res. 48, D882–D889 (2020).

103. Bedrosian, T. A., Quayle, C., Novaresi, N. & Gage, F. H. Early life experience drives structural variation of neural genomes in mice. Science 359, 1395–1399 (2018).

104. Khan, A. et al. JASPAR 2018: update of the open-access database of transcription factor binding profiles and its web framework. Nucleic Acids Res. 46, D260–D266 (2018).

105. Wang, F. et al. RNAscope: a novel in situ RNA analysis platform for formalin-fixed, paraffin-embedded tissues. J. Mol. Diagn. 14, 22–29 (2012).

106. Bodea, L.-G. et al. Neurodegeneration by activation of the microglial complement-phagosome pathway. J. Neurosci. 34, 8546–8556 (2014).

107. Filice, F., Vörckel, K. J., Sungur, A. Ö., Wöhr, M. & Schwaller, B. Reduction in parvalbumin expression not loss of the parvalbumin-expressing GABA interneuron subpopulation in genetic parvalbumin and shank mouse models of autism. Mol. Brain 9, (2016).

108. O’Driscoll, C., Kaufmann, W. E. & Bressler, J. Relationship between Mecp2 and NFκb signaling during neural differentiation of P19 cells. Brain Res. 1490, 35–42 (2013).

